# First estimation of the scale of canonical 5’ splice site GT>GC mutations generating wild-type transcripts and their medical genetic implications

**DOI:** 10.1101/479493

**Authors:** Jin-Huan Lin, Xin-Ying Tang, Arnaud Boulling, Wen-Bin Zou, Emmanuelle Masson, Yann Fichou, Loann Raud, Marlène Le Tertre, Shun-Jiang Deng, Isabelle Berlivet, Chandran Ka, Matthew Mort, Matthew Hayden, Gerald Le Gac, David N. Cooper, Zhao-Shen Li, Claude Férec, Zhuan Liao, Jian-Min Chen

## Abstract

It has long been known that canonical 5’ splice site (5’SS) GT>GC mutations may be compatible with normal splicing. However, to date, the true scale of canonical 5’SS GT>GC mutations generating wild-type transcripts, both in the context of the frequency of such mutations and the level of wild-type transcripts generated from the mutation alleles, remain unknown. Herein, combining data derived from a meta-analysis of 45 informative disease-causing 5’SS GT>GC mutations (from 42 genes) and a cell culture-based full-length gene splicing assay of 103 5’SS GT>GC mutations (from 30 genes), we estimate that ∼15-18% of the canonical GT 5’SSs are capable of generating between 1 and 84% normal transcripts as a consequence of the substitution of GT by GC. We further demonstrate that the canonical 5’SSs whose substitutions of GT by GC generated normal transcripts show stronger complementarity to the 5’ end of U1 snRNA than those sites whose substitutions of GT by GC did not lead to the generation of normal transcripts. We also observed a correlation between the generation of wild-type transcripts and a milder than expected clinical phenotype but found that none of the available splicing prediction tools were able to accurately predict the functional impact of 5’SS GT>GC mutations. Our findings imply that 5’SS GT>GC mutations may not invariably cause human disease but should also help to improve our understanding of the evolutionary processes that accompanied GT>GC subtype switching of U2-type introns in mammals.

## INTRODUCTION

The vast majority of eukaryotic introns are spliced by the U2 spliceosome (the only alternative U12 spliceosome is responsible for <0.5% of all introns [1-3]), which interacts with RNA sequences specifying the 5’ and 3’ splice sites [4, 5]. In vertebrates, the 9-bp consensus sequence for the U2-type 5’ splice site (5’SS) has traditionally been described as 5’-MAG/GURAGU-3’ (where M denotes C or A, R denotes A or G and / denotes the exon-intron boundary; the corresponding nucleotide positions are denoted −3_-1/+1_+6) although in reality this consensus sequence does not reflect the true extent of sequence variability [6-11]. Base-pairing of this 9-bp sequence with 3’-GUCCAUUCA-5’ at the 5’ end of U1 snRNA (Figure 1A) is critical for splicing to occur [10, 12-15]. Although the GT dinucleotide in the first two intronic positions (in the context of DNA sequence) is the most highly conserved portion of the U2-type 5’SS, it was reported, as early as 1983, that GC occasionally occurs in place of GT [16-18]. Subsequent genome-wide analyses have established that this non-canonical 5’SS GC is present as wild-type in ∼1% of human U2-type introns [2, 7, 8, 19, 20]. Importantly, the remaining nucleotides in these evolutionarily fixed non-canonical GC 5’SSs exhibit a stronger complementarity to the 3’-GUCCAUUCA-5’ sequence at the 5’ end of U1 snRNA than those in the canonical GT 5’SSs (Figure 1A), thereby in all likelihood compensating for the decreased complementarity between the 5’SS and the 5’ end of U1 snRNA due to the U to C substitution [7, 8]. Comparative genome analyses have also revealed frequent switching of U2-type introns from the canonical 5’SS GT subtype to the non-canonical 5’SS GC subtype during mammalian evolution [8, 21]. Finally, GC has recently been ranked first among the six non-canonical 5′SSs identified by genome-wide RNA-seq analysis and splicing reporter assays [22].

**Figure 1.**
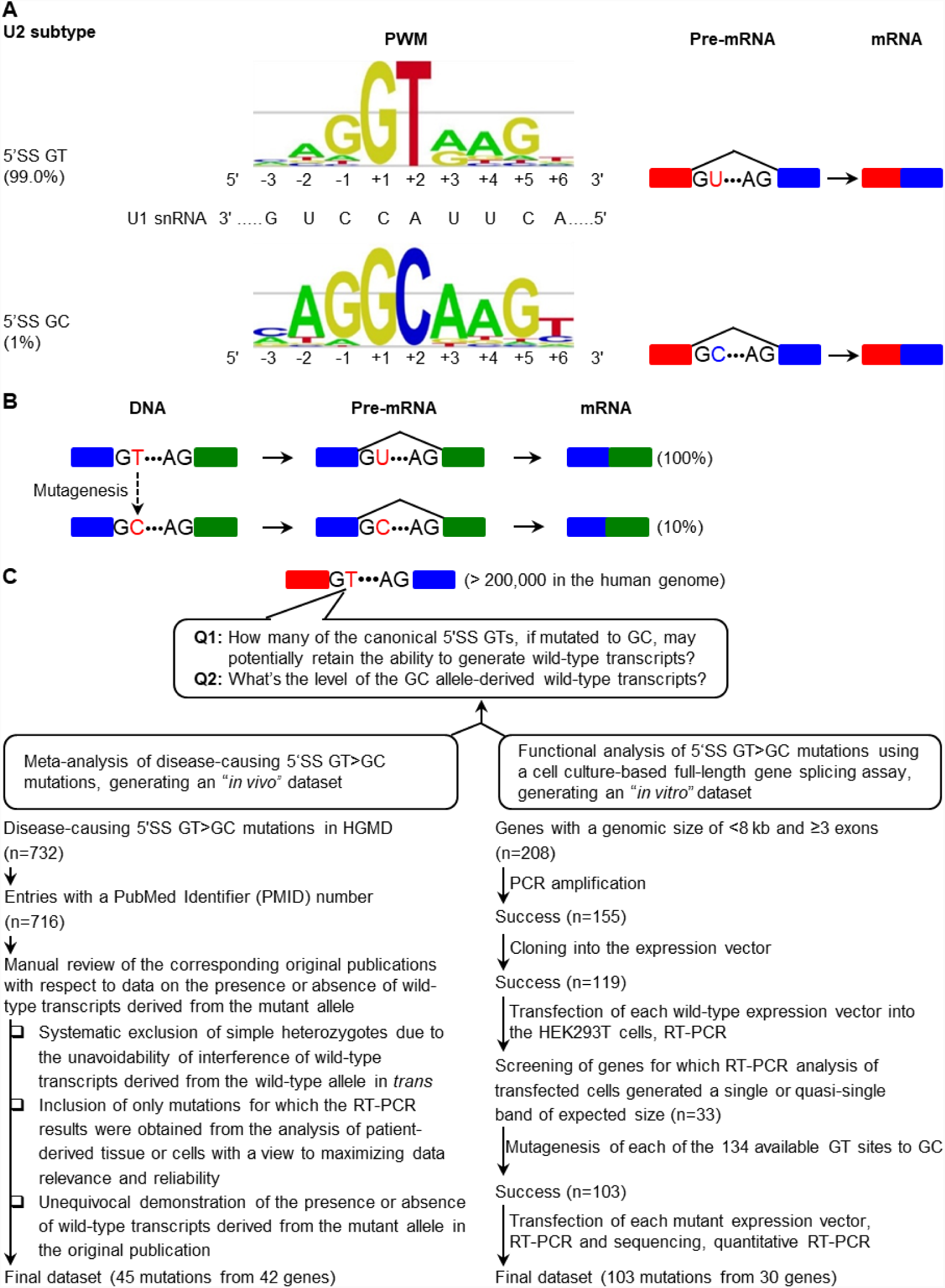
Background Information, Aims and Analytical Strategy of the Study. (A) Current knowledge of the canonical 5’ splice sites (5’SS) GT and non-canonical 5’SS GC in the human genome in terms of their relative abundance of U2-type introns, their corresponding 9-bp 5’SS signal sequence position weight matrices (PWM) and their associated splicing outcomes. The two PWM illustrative figures were taken from Leman et al. Novel diagnostic tool for prediction of variant spliceogenicity derived from a set of 395 combined *in silico/in vitro* studies: an international collaborative effort. Nucleic Acids Res. 2018;46(15):7913-7923 [95] (an Open Access article distributed under the terms of the Creative Commons Attribution Non-Commercial License). (B) Illustration of the first experimental evidence showing that a 5’SS GT>GC mutation may retain the ability to generate wild-type transcripts, albeit at a much reduced level (∼10% of normal in [23, 24]). (C) Aim and analytical strategy of the study.

The finding that GC occasionally occurs instead of GT within the canonical 5’SS in some vertebrate genes implies that substitution of the canonical 5’SS GT by GC (termed a 5’SS GT>GC mutation) may allow normal splicing to occur. The first direct experimental evidence supporting such a postulate came in the late 1980s; analyses of both the splicing products of *in vitro* transcribed rabbit beta globin (*Hbb*) RNA in a HeLa cell nuclear extract and the splicing products of the *Hbb* gene transiently expressed in HeLa cells demonstrated that, of all the possible single nucleotide substitutions of the canonical 5’SS GT of the second and last intron of *Hbb*, only the substitution of T by C was compatible with normal splicing, albeit at a much reduced rate (approximately 10% of normal; see also Figure 1B) [23, 24]. Further supporting evidence came from the study of disease-causing 5’SS GT>GC mutations, some of which were reported to generate wild-type transcripts (see below). Additionally, the activation of cryptic non-canonical 5’SS GC has also been reported as a consequence of some disease-causing mutations [25, 26].

The above notwithstanding, to date, the scale of canonical 5’SS GT>GC mutations generating wild-type transcripts, both in the context of the frequency of such mutations and the level of wild-type transcripts generated by such mutations, remain unknown owing to the intrinsic complexity of splicing [11, 27-29] and the lack of suitable model systems for study. This issue has important implications for medical genetics since mutant genotypes retaining even a small fraction of their normal function may differ significantly from null genotypes in terms of their associated clinical phenotypes (e.g., only 5% of normal *CFTR* gene expression is enough to prevent the lung manifestations of cystic fibrosis [30, 31]). Herein, we attempted to address this issue by employing two distinct but complementary approaches in concert.

## RESULTS AND DISCUSSION

### Estimation by Meta-Analysis of Disease-Causing 5’SS GT>GC Mutations

First, we performed a meta-analysis of disease-causing 5’SS GT>GC mutations logged in the Professional version of Human Gene Mutation Database (HGMD; as of June 2017) [32], with a view to generating an “*in vivo*” dataset to estimate the scale of 5’SS GT>GC mutations generating wild-type transcripts. Employing a stringent approach (Figure 1C), we identified 45 disease-causing 5’SS GT>GC mutations (from 42 genes) that were informative with respect to the presence or absence of wild-type transcripts derived from the mutant allele (Table 1; see Supplementary Table S1 for more information including affected intron, reference mRNA accession number, chromosomal location, hg38 coordinate, and patient-derived tissue or cells used for RT-PCR analysis, etc.). It should be noted that the assignments of “presence” or “absence” of mutant allele-derived wild-type transcripts depended upon the agarose gel evaluation of RT-PCR products as described in the corresponding original publications. Thus, we conservatively annotated an isolated case (i.e., the *PCCB* c.183+2T>C mutation), which was not found to generate wild-type transcripts on agarose gel evaluation of RT-PCR products but was found to generate <0.1% normal wild-type transcripts by means of quantitative RT-PCR [33], as generating no wild-type transcripts.

**Table 1.**
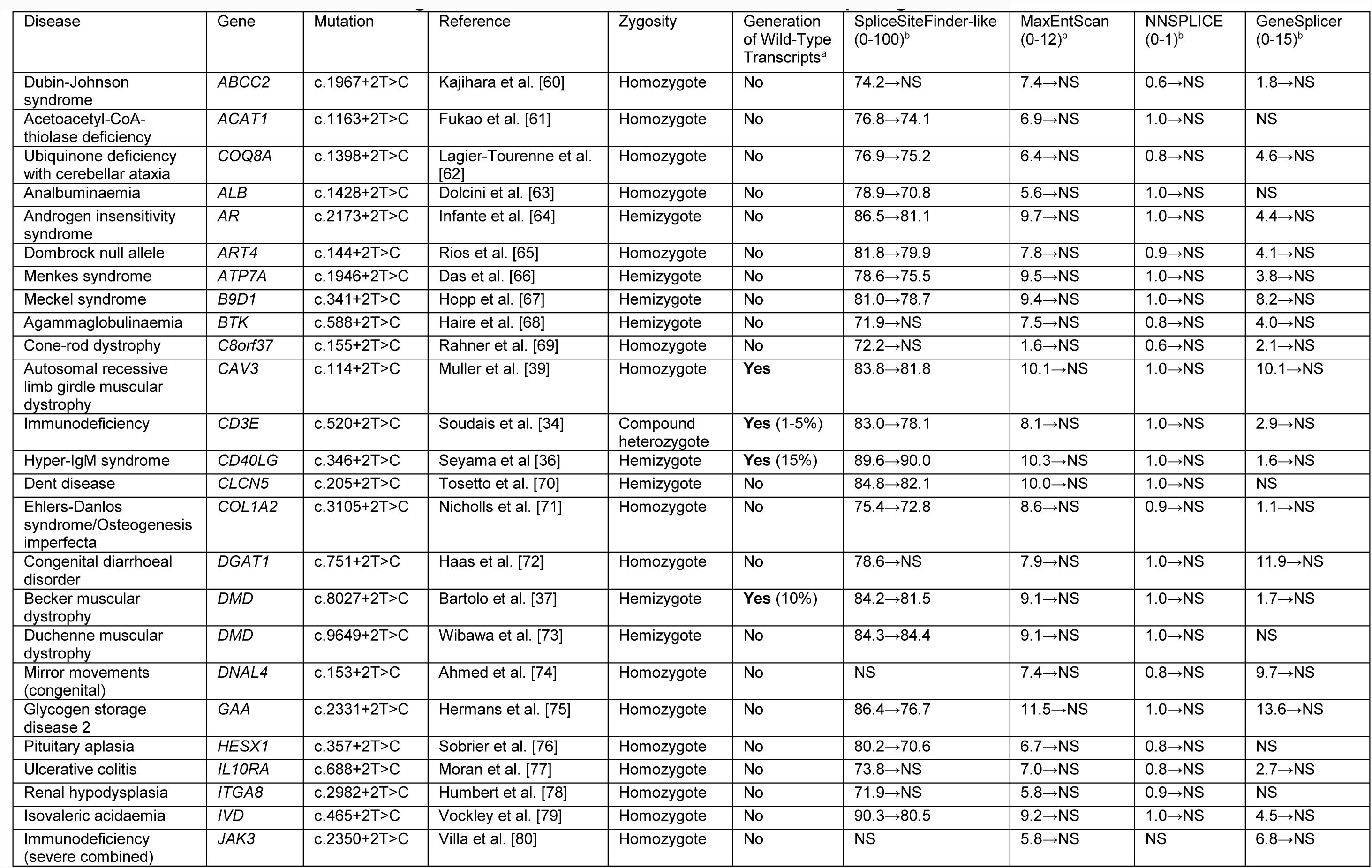
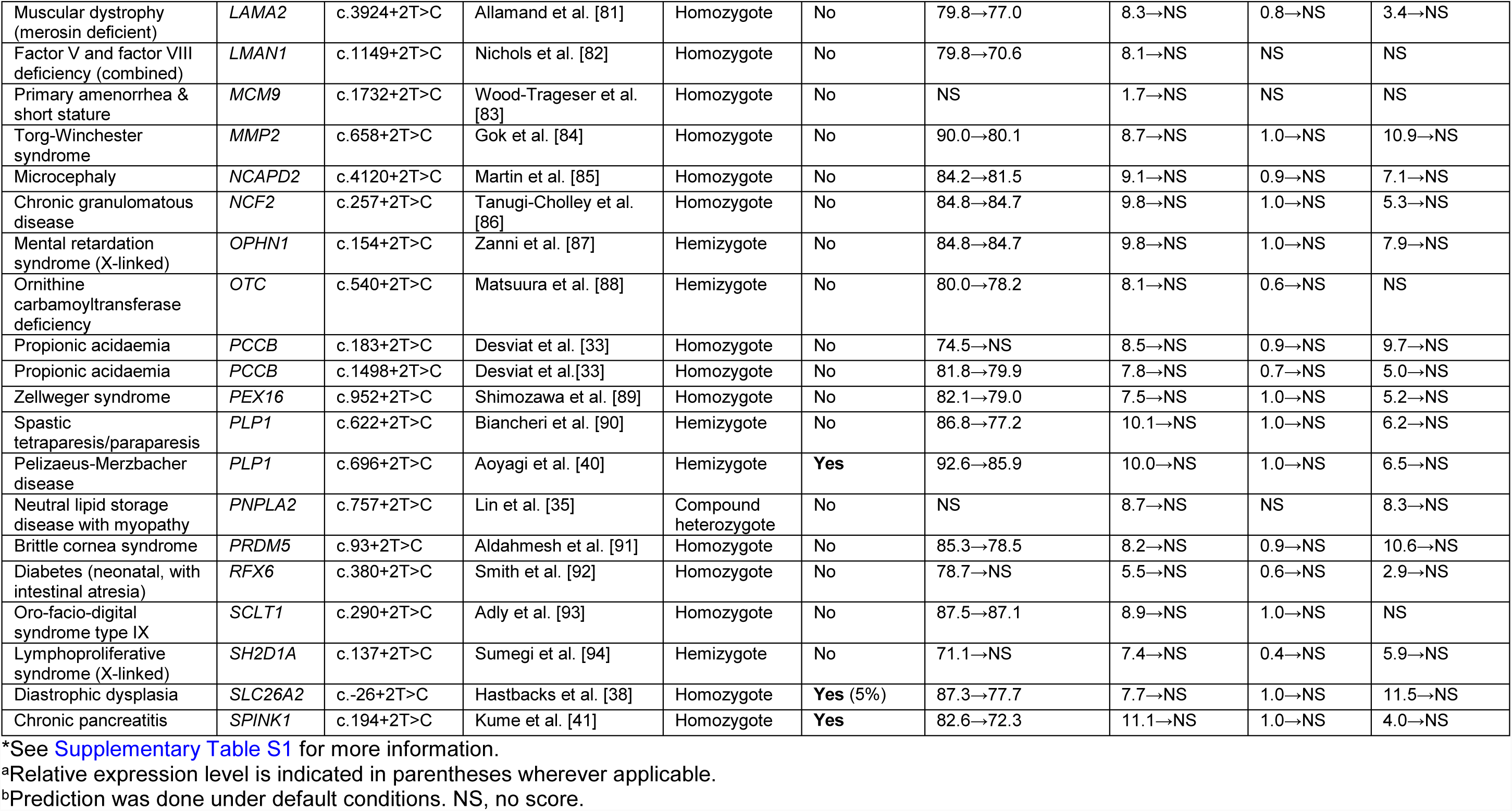
The 45 Informative Disease-Causing 5’SS GT>GC Mutations and their Predicted Splicing Effects*.

The 45 informative 5’SS GT>GC mutations comprised 30 homozygotes, 13 hemizygotes and 2 compound heterozygotes (Table 1). Whilst the presence or absence of wild-type transcripts derived from the mutant allele was straightforward for all homozygous or hemizygous mutations included, the two compound heterozygotes required special treatment. In the case of the *CD3E* c.520+2T>C mutation, the pathogenic *CD3E* mutation in *trans* was a nonsense mutation in exon 6. Sequencing of the patient-derived, normal-sized RT-PCR products failed to demonstrate the exon 6 mutation, suggesting that the wild-type transcripts were derived from the c.520+2T>C allele [34]. In the case of the *PNPLA2* c.757+2T>C mutation, the second *PNPLA2* mutation in *trans* was a missense mutation, c.749A>C (p.Gln250Pro). RT-PCR analysis detected only the c.749A>C mutant mRNA in skeletal muscle of the patient, indicating the absence of detectable wild-type transcript emanating from the c.757+2T>C allele [35].

15.6% (n=7) of the 45 informative mutations were found to have been capable of generating some correctly spliced transcripts (Table 1). Information on the expression level of the mutant allele-derived wild-type transcripts relative to that of the wild-type transcripts from a normal control (by definition, 100%) was available from four of the seven original publications (i.e., *CD3E* c.520+2T>C [34], *CD40LG* c.346+2T>C [36], *DMD* c.8027+2T>C [37] and *SLC26A2* c.-26+2T>C [38]), which ranged from 1-15% of normal in individual cases (Table 1). All three of the remaining mutations generated both wild-type and aberrant transcripts (i.e., *CAV3* c.114+2T>C [39], *PLP1* c.696+2T>C [40] and *SPINK1* c.194+2T>C [41]); based upon visual inspection of the original gel photographs, the relative expression level of the mutant allele-derived wild-type transcripts in these three cases could also be estimated to fall within the 1-15% range.

Taken together, the meta-analysis of disease-causing mutations suggests that 15.6% of 5’SS GT>GC mutations retained the ability to generate between 1 and 15% correctly spliced transcripts relative to their wild-type counterparts.

### Estimation from the Cell Culture-Based Full-Length Gene Splicing Assay of 5’SS GT>GC Mutations

To corroborate the findings derived from the above “*in vivo*” dataset, we sought to generate an “*in vitro*” dataset of 5’SS GT>GC mutations. In this regard, we have previously used a cell culture-based full-length gene splicing assay to analyze a series of *SPINK1* intronic variants including a 5’SS GT>GC mutation, c.194+2T>C [42, 43]. Specifically, the full-length 7-kb *SPINK1* genomic sequence (including all four exons plus all three introns of the gene) was cloned into the pcDNA3.1/V5-His-TOPO vector [44]. The full-length gene splicing assay preserves better the natural genomic context of the studied mutations as compared to the commonly used minigene splicing assay, a point of importance given the highly context-dependent and combinatorial nature of alternative splicing regulation [45]. Moreover, the full-length gene splicing assay can be readily used to evaluate all intronic variants including those located near the first or last exons of the gene. Despite these advantages, the full-length gene assay cannot easily be applied to large-size genes owing to the technical difficulties inherent in amplifying and cloning long DNA fragments into the expression vector. Finally, it is pertinent to point out that, to functionally evaluate the impact on splicing of any given gene mutation in a transient expression system, it is highly desirable to use of cells of pathophysiological relevance owing to the tissue specificity of the splicing process in some instances [11, 27-29]. However, this may not always be possible in practice, particularly if variants in multiple genes are to be analyzed in large-scale studies. For example, a recent study that measured 5’SS activity in the context of three minigenes was performed in transfected HeLa cells [11]. In the present study, we used HEK293T cells for transfection as previously described [42, 46].

Bearing in mind the aforementioned advantages and disadvantages, we employed a cell culture-based full-length gene splicing assay (Figure 1C). In brief, for various technical and practical reasons, we firstly selected genes whose genomic sizes did not exceed 8 kb (from the translation initiation codon to the translation termination codon) and whose exons numbered ≥3 in order to construct full-length gene expression vectors; we then screened genes, which had yielded a single or quasi-single band of expected size by means of RT-PCR analysis of transfected cells, for subsequent mutagenesis of all available 5’SS GT dinucleotides in the construct (for details on the selected and screened genes, see Supplementary Table S2). In the end, we succeeded in functionally analyzing 103 GT>GC mutations from 30 different genes (Supplementary Table S3). 18.4% (n=19) of these artificially introduced 5’SS GT>GC mutations generated wild-type transcripts (all confirmed by Sanger sequencing; Figure 2 and Supplementary Figure S1), a finding that concurs with the 15.6% value obtained from the meta-analysis of disease-causing 5’SS GT>GC mutations.

**Figure 2.**
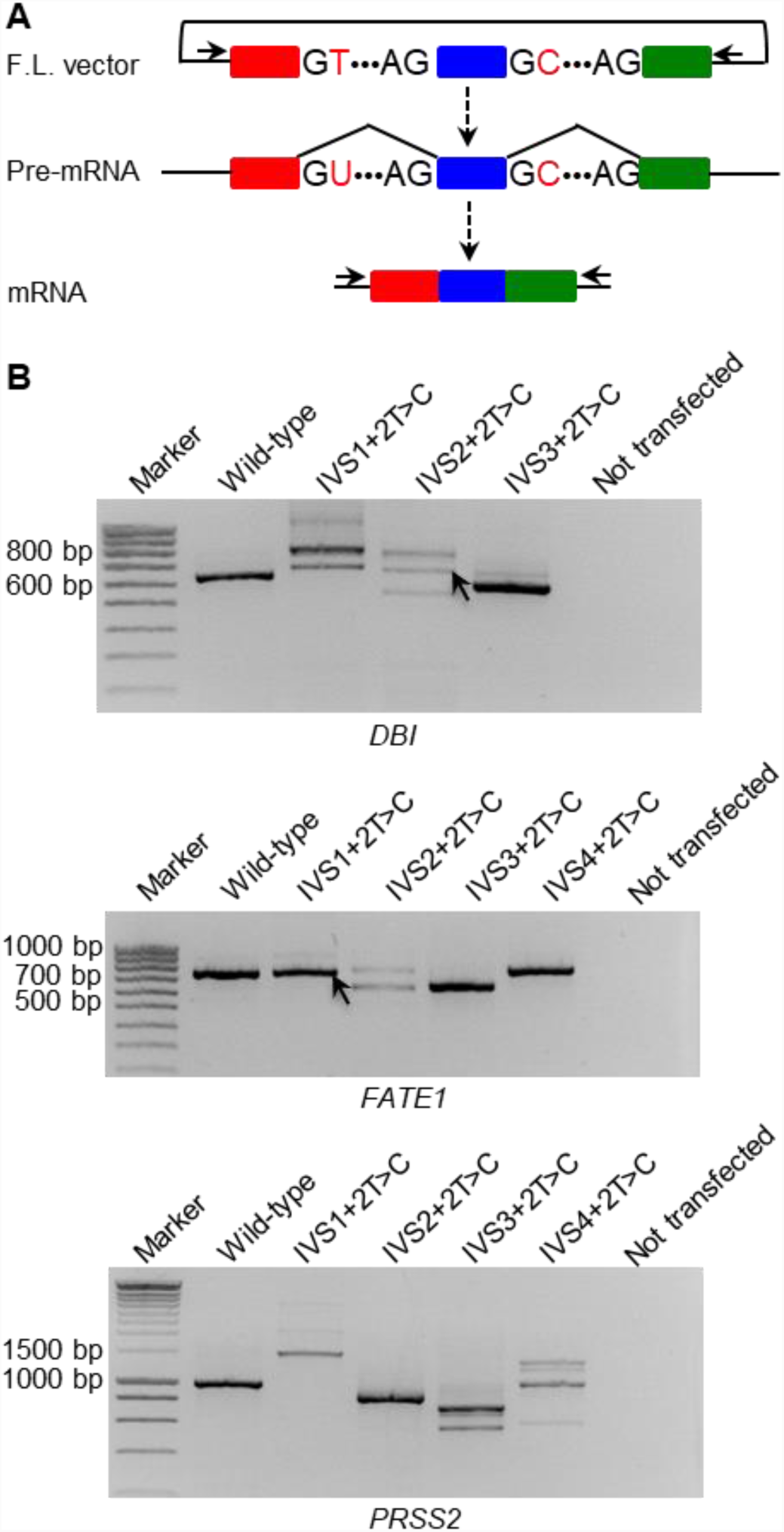
Qualitative Analysis of 5’SS GT>GC Mutations. (A) Illustration of the cell culture-based full-length gene splicing assay in the context of a 5’SS GT>GC mutation generating some wild-type transcripts. The two horizontal arrows indicate the primers (both located within the vector sequence) used to amplify normally spliced transcripts (and also aberrantly spliced transcripts). F.L., full-length. (B) RT-PCR analyses of HEK293T cells transfected with full-length *DBI, FATE1* and *PRSS2* gene expression constructs carrying respectively the wild-type and 5’SS GT>GC mutations as examples. Normal transcripts (confirmed by sequencing) resulting from two of the mutations are indicated by arrows. IVS, InterVening Sequence (i.e., an intron). See Supplementary Figure S1 for all 103 functionally analyzed 5’SS GT>GC mutations.

Only wild-type transcripts were observed for 10 of the aforementioned 19 5’SS GT>GC mutations (e.g., *FATE1* IVS1+2T>C in Figure 2B). In other words, no aberrantly spliced transcripts were observed in these 10 cases. It is possible that aberrantly spliced transcripts may be rendered invisible by RNA degradation mechanisms such as nonsense-mediated mRNA decay (NMD) [47, 48]. One way to test such a possibility is to add an NMD inhibitor such as cycloheximide [49] to the cell culture medium, although this is beyond the scope of the present study. We quantified the relative level of correctly spliced transcripts for these 10 5’SS GT>GC mutations by means of our previously described quantitative RT-PCR method [46, 50, 51]. Here it is pertinent to mention that a co-transfected minigene construct was used as an internal control in this analysis (Figure 3A), a prerequisite to obtain accurate results. As shown in Figure 3B, the relative level of correctly spliced transcripts emanating from these 10 mutations is remarkably similar to that observed for the disease-causing 5’SS GT>GC mutations in terms of the lowest extreme (2-5% vs. 1-5%); however, the functionally obtained highest level of correctly spliced transcripts (84%) is much higher than the corresponding 15% value observed for the disease-causing 5’SS GT>GC mutations (Table 1). We were initially puzzled by this disparity, but this could be accounted for by two considerations. On the one hand, the currently analyzed disease-causing mutations were likely to be biased toward those that generated either no wild-type transcripts or only a low level. On the other hand, given (i) that 5’SS GC may occur as wild-type in the human genome, (ii) the highly degenerate nature of the 5’SS splice signal sequences and (iii) the complex regulation of the splicing process *in vivo*, it is entirely possible that a 5’SS GT>GC mutation may behave similarly to its original wild-type sequence. This notwithstanding, no single GC mutation was noted to have an identical or higher normal splicing activity than its 5’SS GT counterpart (Figure 3B).

**Figure 3.**
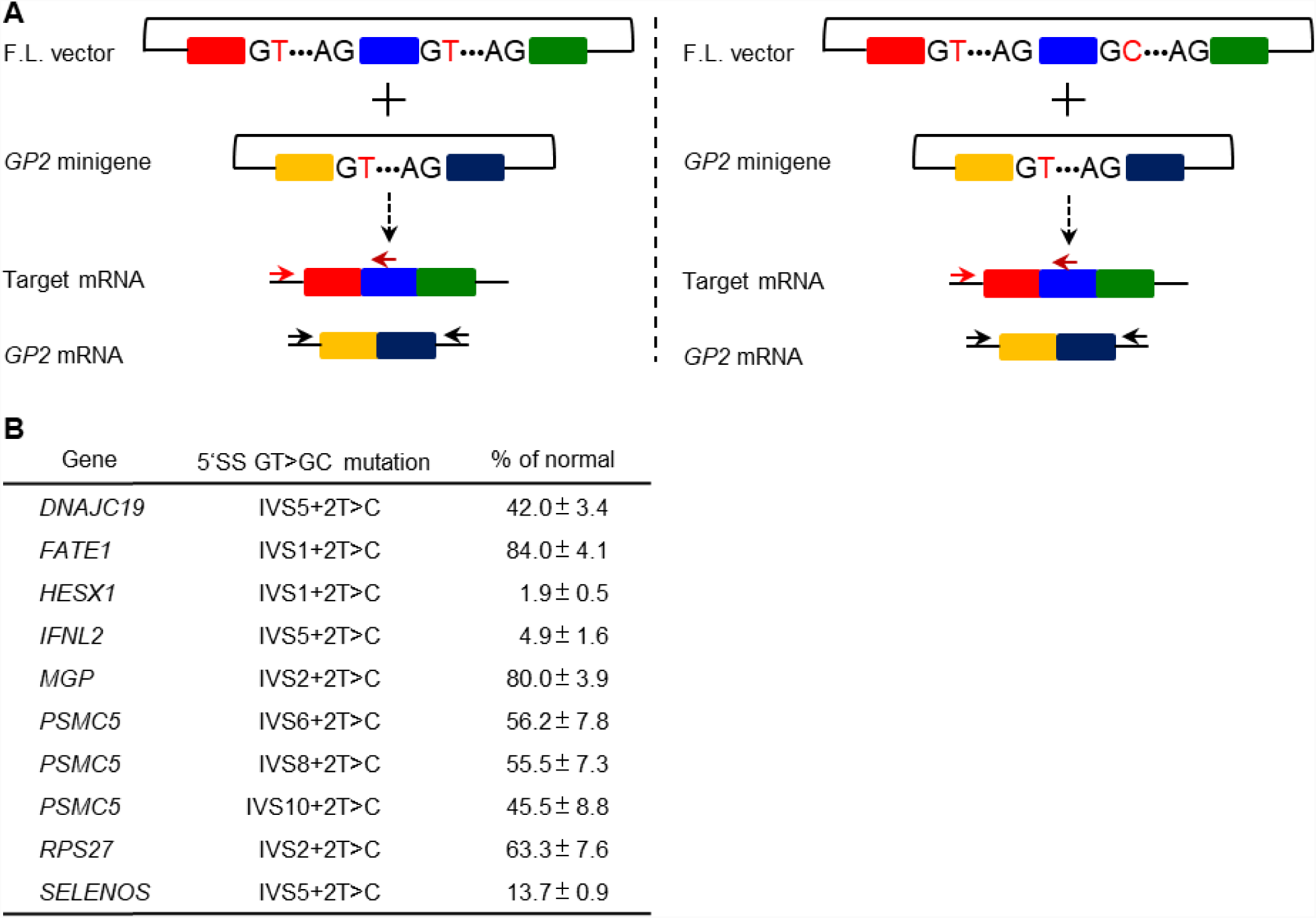
Quantitative Analysis Pertaining to the Relative Level of 5’SS GT>GC Mutation-Derived Wild-Type Transcripts. (**A**) Illustration of one key feature of the quantitative RT-PCR analysis: co-transfection of a minigene expression vector with respectively the full-length wild-type target gene expression vector and the full-length variant target gene expression vector. The minigene was constructed in pGL3 [44] whereas the target gene was constructed in either pcDNA3.1/V5-His-TOPO vector or pcDNA3.1(+). The minigene was used as an internal control for quantifying the expression level of wild-type transcripts generated from either the wild-type or variant target full-length gene. The horizontal arrows indicate the relative positions of the primers used for this purpose. Note that for amplifying the target gene sequence, either a primer pair comprising a forward vector-specific primer and a reverse gene-specific primer (as illustrated) or alternatively a primer pair comprising a forward gene-specific primer and a reverse vector-specific primer was used. This assay was performed exclusively for the 10 5’SS GT>GC mutations that generated only wild-type transcripts. F.L., full-length. (B) Quantitative RT-PCR-determined expression level of the mutant allele-derived correctly spliced transcripts relative to that derived from the corresponding wild-type allele (defined as 100%) in the 10 5’SS GT>GC mutations that generated only wild-type transcripts. Results were expressed as means ± SD from three independent transfection experiments.

Additionally, the single RT-PCR band of wild-type transcript size from either the wild-type *CCDC103* gene or the *CCDC103* IVS1+2T>C mutant (refer to Supplementary Figure S1) was revealed by Sanger sequencing to comprise the correctly spliced transcript and an alternatively spliced transcript; the level of the correctly spliced transcripts generated from the mutant allele was estimated to be ∼18% of that generated from the wild-type allele based upon evaluation of the corresponding sequence peak heights (Supplementary Figure S2). By contrast, we did not attempt to quantify the relative expression level of correctly spliced transcripts for the remaining 8 GT>GC mutations due to the co-presence of aberrantly spliced transcripts (e.g., *DBI* IVS2+2T>C in Figure 2B). Nonetheless, based upon the relative intensities of the wild-type and aberrant transcript bands (Figure 2; Supplementary Figure S1), we consider it unlikely that the relative expression level of correctly spliced transcripts in these cases will have fallen outside of the above experimentally obtained 2-84% range.

Finally, we sequenced some aberrantly spliced transcripts (n=12), which resulted from exon skipping, retention of intronic sequence or deletion of partial exonic sequences (Table 2). Notably, the *PRSS2* IVS4+2T>C mutation activated a cryptic 5’SS GC that is located 15 bp upstream of the normal one, resulting in the deletion of the last 17 bp of exon 4 (i.e., the major band generated by *PRSS2* IVS4+2T>C; Figure 2B).

**Table 2.** Nature of the Sequenced 12 Aberrantly Spliced Transcripts*.

| Gene | Reference mRNA accession number | Mutation | Aberrant Transcripts |
| --- | --- | --- | --- |
| *DBI* | NM_001079862.2 | IVS1+2T>C | 1. Activation of a cryptic 5'SS GT which is located 152 bp downstream of the normal one, resulting in the retention of the first 154 bp of the intron 1 sequence.<br>2. Activation of a cryptic 5'SS GT which is located 28 bp downstream of the normal one, resulting in the retention of the first 30 bp of the intron 1 sequence. |
|  |  | IVS3+2T>C | Exon 3 skipping |
| *FABP7* | NM_001446.4 | IVS1+2T>C | Activation of a cryptic 5'SS GT which is located 2 bp downstream of the normal one, resulting in the retention of the first 4 bp of the intron 1 sequence. |
|  |  | IVS2+2T>C | Activation of a cryptic 5'SS GT which is located 3 bp upstream of the normal one, resulting in the deletion of the last 5 bp of exon 1. |
| *HESX1* | NM_003865.2 | IVS2+2T>C | Exon 2 skipping |
|  |  | IVS3+2T>C | Exon 3 skipping |
| *IL10* | NM_000572.3 | IVS1+2T>C | Activation of a cryptic 5'SS GT which is located 2 bp downstream of the normal one, resulting in the retention of the first 4 bp of the intron 1 sequence. |
|  |  | IVS4+2T>C | Activation of a cryptic 5'SS GT which is located 19 bp upstream of the normal one, resulting in the deletion of the last 21 bp of exon 4. |
| *PRSS2* | NM_002770.3 | IVS4+2T>C | Activation of a cryptic 5'SS GC which is located 15 bp upstream of the normal one, resulting in the deletion of the last 17 bp of exon 4. |
| *SPINK1* | NM_003122.3 | IVS1+2T>C | Activation of a cryptic 5'SS GT which is located 138 bp downstream of the normal one, resulting in the retention of the first 140 bp of the intron 1 sequence. |
| *UQCRB* | NM_006294.4 | IVS1+2T>C | Activation of a cryptic 5'SS GT which is located 10 bp upstream of the normal one, resulting in the deletion of the last 12 bp of exon 1. |

\*See Supplementary Figure S1 for the corresponding RT-PCR products

### Integrated Estimation from the Two Distinct but Complementary Datasets

We obtained remarkably similar findings in terms of both the frequency of 5’SS GT>GC mutations generating wild-type transcripts and the lowest relative level of mutant allele-derived wild-type transcripts from two quite distinct yet complementary datasets. The consistently lowest relative level of mutant allele-derived wild-type transcripts across the two datasets suggested that the gel-based analytical method is sensitive enough to detect as little as ∼1% of normally spliced transcripts. The apparent disparity in terms of the highest relative level of mutant-derived wild-type transcripts between the two datasets can however be accounted for largely by the selection bias inherent to disease-causing mutations. Therefore, we estimate that some 15-18% of 5’SS GT>GC mutations generate between 1 and 84% of wild-type transcripts.

### Exploration of the Mechanisms Underlying the Generation or Not of Wild-Type Transcripts by 5’SS GT>GC Mutations

As mentioned above, canonical GT and non-canonical GC 5’SSs in the human genome exhibit different patterns of sequence conservation, the latter showing stronger complementarity to the 3’-GUCCAUUCA-5’ sequence at the 5’ end of U1 snRNA (Figure 1A). We surmised that the canonical 5’SSs whose substitutions of GT by GC generated normal transcripts (termed group 1) should also exhibit stronger complementarity to the aforementioned 9-bp sequence than those sites whose substitutions of GT by GC did not lead to the generation of normal transcripts (termed group 2). We therefore extracted the 9-bp sequence tracts surrounding the corresponding groups of the 45 disease-causing 5’SS GT>GC mutations (Supplementary Tables S1) and those of the 103 functionally analyzed 5’SS GT>GC mutations (Supplementary S3). Comparison of the resulting pictograms confirmed our postulate in both contexts, the respective pictograms for the combined group 1 mutations (n=26) and combined group 2 mutations (n=122) being provided in Figure 4. It should be emphasized that the surrounding 9-bp sequence tract is an important (but certainly not the only) factor in determining whether or not a given 5’SS GT>GC mutation will generate some wild-type transcripts. A simple example may be used to illustrate this point: the *DMD* c.8027+2T>C mutation (which generates 10% of wild-type transcripts) contrasts with the *NCAPD2* c.4120+2T>C mutation (which generates no wild-type transcripts) despite occurring in an identical 9-bp sequence tract, AAGGTATGA (see Supplementary Table S1).

**Figure 4.**
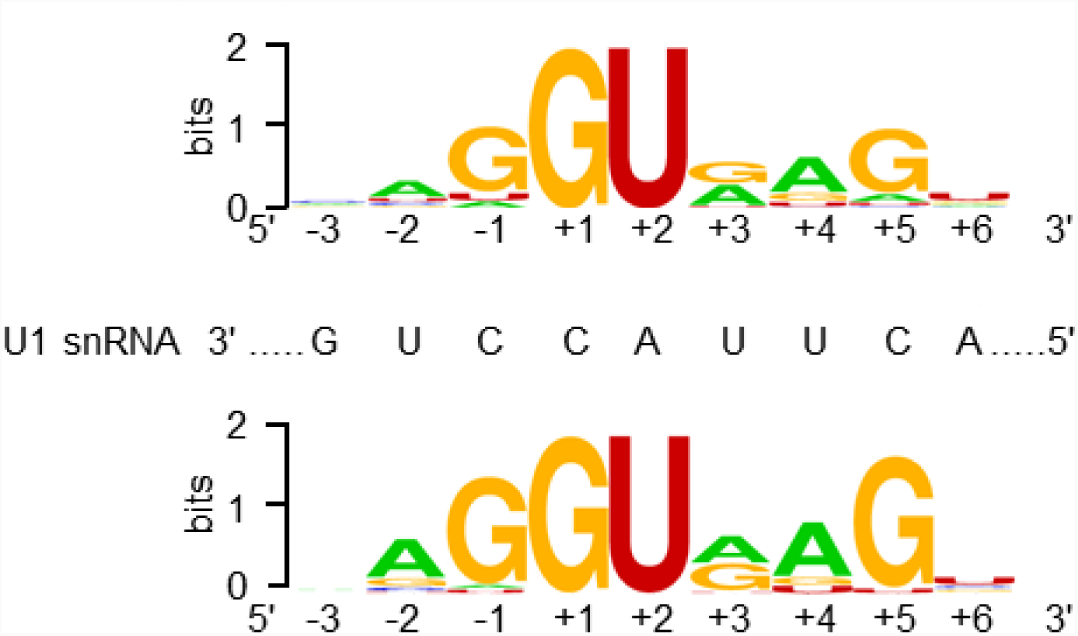
Pictogram Analysis of the 5’SSs Under Study. Comparison of the pictogram of the 122 5’SSs whose substitutions of GT by GC did not lead to the generation of normal transcripts (upper panel) and that of the 26 5’SSs whose substitutions of GT by GC generated normal transcripts (lower panel). Middle panel shows the 5’ end sequence of U1 snRNA that is complementary to the 9-bp U2-type 5’SS signal sequence. 5’SS signal sequences are shown as RNA sequence.

We also explored whether the creation or disruption of splice enhancer/silencer motifs by the 5’SS GT>GC mutations could be associated with the generation or not of some wild-type transcripts. To this end, we employed ESEfinder and RESUE-ESE provided by the Alamut suite under default conditions. We were unable to draw any meaningful conclusions, primarily due to the short and degenerate nature of the splicing enhancer/silencer binding motifs.

### Correlation Between the Retention of Wild-Type Transcripts and a Milder Than Expected Phenotype

Given that even the retention of a small fraction of normal gene function may significantly impact the clinical phenotype, we reviewed the original publications describing the seven disease-causing 5’SS GT>GC mutations that generated at least some wild-type transcript (Table 1) with respect to the accompanying genotypic and phenotypic descriptions. In six cases, the mutations were specifically described as being associated with mild clinical phenotypes as compared to their classical disease counterparts (see Supplementary Table S1). In the remaining case (*SPINK1* c.194+2T>C), the original publication [41] was not informative in this regard; however, it is known that homozygosity for this mutation causes chronic pancreatitis with variable expressivity [52] whereas null *SPINK1* genotypes cause severe infantile isolated exocrine pancreatic insufficiency [53].

The above correlation between the retention of some wild-type transcripts and a milder than expected phenotype prompted us to postulate that 5’SS GT>GC mutations previously reported to confer a milder than expected phenotype but having no supportive patient-derived transcript expression data, may be collectively associated with a non-canonical 5’SS GC signal. We collated a total of six such mutations (i.e., *CYB5R3* c.463+2T>C [54], *HBB* c.315+2T>C [55], *HPRT* c.485+2T>C [56], *LAMB2* c.3327+2T>C [57], *LMNA* c.1968+2T>C [58] and *MTTP* c.61+2T>C [59]; Supplementary Table S4). In this regard, two points require clarification. First, in two cases, patient-derived transcript expression data were available [56, 57]; these cases were however addressed here because the corresponding expression data were insufficiently informative for them to be listed in Supplementary Table S1 (for explanations, see Supplementary Table S4). Second, five of these six mutations (all germline) were derived from the HGMD dataset whereas the remaining one (*LMNA* c.1968+2T>C) [58], a somatic mutation, was obtained from a literature search; this somatic mutation was included owing to its clear phenotypic impact. Pictogram analysis of the six corresponding 9-bp canonical 5’SSs did reveal a non-canonical 5’SS GC signal (Supplementary Figure S3). Notably, one of the mutations affected the splice donor splice site of *HBB* intron 2 (i.e., *HBB* c.315+2T>C) [55], site of the previously analyzed orthologous mutation in the rabbit *Hbb* gene [23, 24]. We were able to study the effect of the *HBB* intron 2 GT>GC mutation on splicing by means of the full-length gene assay and found that it had indeed retained the ability to generate normal *HBB* transcripts (Figure 5).

**Figure 5.**
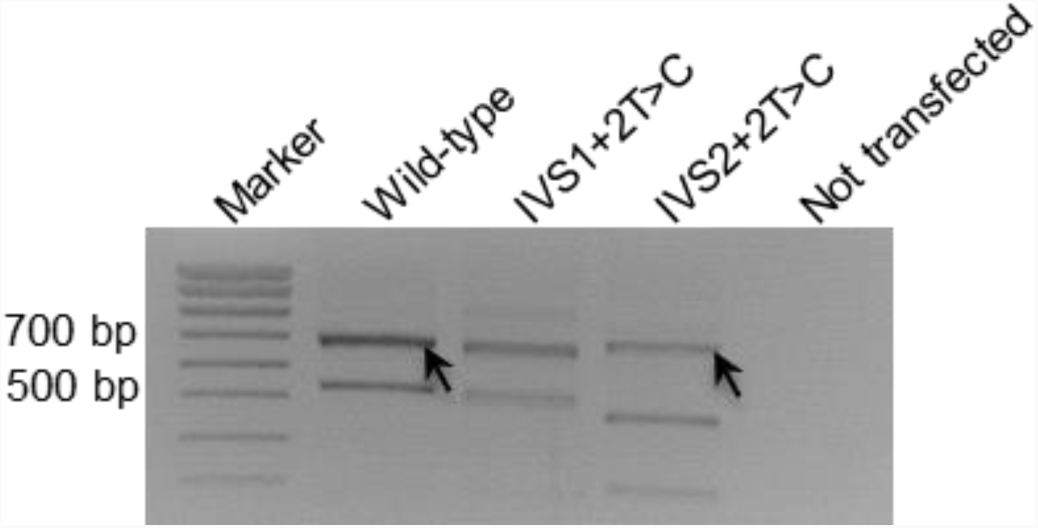
Functional Characterization of the *HBB* c.315+2T>C mutation. RT-PCR analyses of HEK293T cells transfected with full-length *HBB* gene expression constructs carrying respectively the wild-type and two 5’SS GT>GC mutations. Wild-type transcripts (confirmed by sequencing) resulting from the wild-type and the IVS2+2T>C (i.e., c.315+2T>C) mutation are indicated by arrows. The *HBB* c.315+2T>C mutation was previously reported to be associated with a mild phenotype [55]. IVS, InterVening Sequence (i.e., an intron).

### Prediction of the Functional Effect of 5’SS GT>GC Mutations

Finally, it is important to point out that none of the splicing prediction tools were able to accurately predict the functional effect of 5’SS GT>GC mutations. For example, we analyzed the 45 disease-causing 5’SS GT>GC mutations as well as the 19 functionally analyzed 5’SS GT>GC mutations that generated some wild-type transcripts by means of the widely used Alamut^®^ software suite under default conditions. Whereas SpliceSiteFinder-like tended to predict a slightly reduced score, MaxEntScan, NNSPLICE and GeneSplicer invariably gave no scores, for all mutations tested (Table 1; Supplementary Table S3).

### Conclusions

Based upon complementary data from the meta-analysis of 45 disease-causing 5’SS GT>GC mutations and the cell culture-based full-length gene splicing analysis of 103 5’SS GT>GC mutations, we have provided a first estimate of ∼15-18% for the proportion of canonical GT 5’SSs that are capable of generating between 1 and 84% normal transcripts in case of the substitution of GT by GC. Extrapolation of the 15-18% value to the entire human genome implies that in at least 30,000 U2-type introns, the substitution of 5’SS GT by GC would result in the retention of partial ability to generate wild-type transcripts. Given that even the retention of 5% normal transcripts can significantly ameliorate a patient’s clinical phenotype, our findings imply the potential existence of hundreds or even thousands of disease-causing 5’SS GT>GC mutations that may underlie relatively mild clinical phenotypes. Given that 5’SS GT>GC mutations can also give rise to relatively high levels of wild-type transcripts, our findings imply that 5’SS GT>GC mutations may not invariably cause human disease. Apart from their direct implications for medical genetics, our findings may also help to improve our understanding of the evolutionary processes that accompanied the GT>GC subtype switching of U2-type introns in mammals [8, 21].

## MATERIALS AND METHODS

### Meta-Analysis of Disease-Causing 5’SS GT>GC Mutations

Human disease-causing 5’SS GT>GC mutations logged in the Professional version of the Human Gene Mutation Database (HGMD; http://www.hgmd.cf.ac.uk/ac/index.php; as of June 2017) [32] were used as starting material. The procedure of the meta-analysis is described in Figure 1C.

### Cell Culture-Based Full-Length Gene Splicing Assay

Outline of the cell culture-based full-length gene splicing assay is illustrated in Figure 1C.

#### Amplification of full-length gene sequences

For this experiment, we focused on genes whose genomic sizes were <8 kb (from the translation initiation codon to the translation termination codon) and whose exons numbered ≥3. Long-range PCR was performed in a 25 µL reaction mixture containing 0.5 U KAPA HiFi HotStart DNA Polymerase (Kapa Biosystems), 0.75 µL KAPA dNTP Mix (300 µM final), 5 µL 5 × KAPA HiFi Buffer, 50 ng DNA, and 0.3 µM forward and reverse primers (primer sequences available upon request). The PCR program comprised an initial denaturation at 95°C for 5 min, followed by 30 cycles of denaturation at 98°C for 20 s, annealing at 66°C for 15 s, extension at 72°C for 1 min/kb, and a final extension at 72°C for 5 min. In some of the cases where the desired fragments could not be obtained, a second amplification was attempted: PCR was performed using 50 ng DNA in a 50 µL reaction mixture with 2.5 U TaKaRa LA Taq DNA polymerase (TaKaRa), 8 µL dNTP Mixture (400 µM final), 5 µL 10 × LA PCR Buffer, and 1 µM forward and reverse primers; thermal cycling conditions were initial denaturation at 94°C for 1 min, 30 cycles of denaturation at 98°C for 10 s, annealing and extension at 68°C for 1 min/kb, and a final extension at 72°C for 10 min.

#### Cloning of the amplified full-length wild-type gene sequences into the expression vector

Early experiments were performed by means of TA cloning. In those cases in which the PCR products contained multiple bands, the band of the expected size was gel purified using the QIAquick Gel Extraction Kit (Qiagen) and 3’-A overhangs added; in cases where a single and expected band was obtained, 3’-A overhangs were directly added to the PCR products amplified from the KAPA HiFi HotStart DNA Polymerase (this step was omitted for those amplified using the TaKaRa LA Taq DNA polymerase). The resulting products were cloned into the pcDNA3.1/V5-His-TOPO vector (Invitrogen) in accordance with the manufacturer’s instructions. Transformation was performed using Stellar Competent Cells (TaKaRa) or XL10-Gold Ultracompetent Cells (Agilent Technologies). Transformed cells were spread onto LB agar plates with 50 µg/mL ampicillin and incubated at 37°C overnight. Plasmid constructs containing inserts in the right orientation were selected by PCR screening using the HotStarTaq Master Mix Kit (Qiagen).

Later experiments were performed by means of in-fusion cloning. PCR products of the expected size were purified using the QIAquick Gel Extraction Kit (Qiagen) after gel electrophoresis. The purified products were cloned into *EcoR*I restriction site of the linearized pcDNA3.1(+) vector with the In-Fusion HD Cloning kit (TaKaRa) according to the manufacturer’s instructions. Transformation was performed using Stellar Competent Cells (TaKaRa) or XL10-Gold Ultracompetent Cells (Agilent Technologies). Transformed cells were spread onto LB agar plates with 50 µg/mL ampicillin and incubated at 37°C overnight. Plasmid constructs containing inserts were confirmed by PCR using the HotStarTaq Master Mix Kit (Qiagen).

#### Mutagenesis

Variants were introduced into the wild-type full-length gene expression constructs by means of the QuikChange II XL Site-Directed Mutagenesis Kit (Agilent Technologies). Mutagenesis was performed in a 25.5 μL mixture containing 1.25 U PfuUltra HF DNA polymerase, 0.5 μL dNTP mix, 2.5 μL 10× reaction buffer, 1.5 μL QuikSolution, 100 ng wild-type plasmid, and 62.5 ng each mutagenesis primer (primer sequences available upon request). The PCR program had an initial denaturation at 95°C for 2 min, followed by 18 cycles of denaturation at 95°C for 1 min, annealing at 60°C for 50 s, and extension at 68°C for 1 min/kb, and a final extension at 68°C for 7 min. The PCR products were transformed into XL10-Gold Ultracompetent cells (Agilent Technologies) after treated with *Dpn*I at 37°C for 1 h. Transformed cells were spread onto LB agar plates with 50 µg/mL ampicillin and incubated at 37°C overnight. Selected colonies were cultured overnight. Plasmids were isolated using the QIAprep Spin Miniprep Kit (Qiagen) and the successful introduction of the desired mutations was validated by DNA sequencing with the BigDye Terminator v1.1 Cycle Sequencing Kit (Applied Biosystems).

#### Cell culture, transfection, RNA extraction, and reverse transcription

Human embryonic kidney 293T (HEK293T) cells were cultured in the Dulbecco’s modified Eagle’s medium (BioWhittaker) with 10% fetal calf serum (Eurobio). 3.5 × 10^5^ cells were seeded per well in 6-well plates 24 h before transfection. For conventional RT-PCR analyses, 1 µg wild-type or variant plasmid, mixed with 2 µL jetPEI DNA transfection reagent (Polyplus-transfection), was used for transfection per well. For real-time quantitative RT-PCR analyses, 500 ng wild-type or variant plasmid was mixed with 500 ng pGL3-GP2 minigene for transfection [44, 46, 50]. Forty-eight hours after transfection, total RNA was extracted using the RNeasy Mini Kit (Qiagen). RT was performed with 200 U SuperScript III Reverse Transcriptase (Invitrogen), 500 µM dNTPs, 4 µL 5 × First-Strand Buffer, 5 mM dithiothreitol, 2.5 µM 20mer-oligo (dT), and 1 µg total RNA. The resulting complementary DNA (cDNA) were treated with 2U RNaseH (Invitrogen) to degrade the remaining RNA.

#### Conventional RT-PCR analyses and sequencing of the resulting products

Conventional RT-PCR was performed in a 25-μL reaction mixture containing 12.5 μL HotStarTaq Master Mix (Qiagen), 1 μL cDNA, and 0.4 μM each primer (5’-GGAGACCCAAGCTGGCTAGT-3’ (forward) and 5’-AGACCGAGGAGAGGGTTAGG-3’ (reverse) for TA cloning-obtained plasmids (both primers are located within the pcDNA3.1/V5-His-TOPO vector sequence); 5’-TAATACGACTCACTATAGGG-3’ (forward) and 5’-TAGAAGGCACAGTCGAGG-3’ (reverse) for in-fusion cloning-obtained plasmids (both primers are located within the pcDNA3.1(+) vector sequence)). The PCR program had an initial denaturation step at 95°C for 15 min, followed by 30 cycles of denaturation at 94°C for 45 s, annealing at 58°C for 45 s, and extension at 72°C for 1 min/kb (in the step to screen wild-type genes for which RT-PCR analysis of transfected cells generated a single or quasi-single band of expected size) or for 2 min (in the step to analyze the splicing outcomes of 5’SS GT>GC mutations), and a final extension step at 72°C for 10 min. RT-PCR products of a single band were cleaned by ExoSAP-IT (Affymetrix). In the case of multiple bands, the band corresponding to the normal-sized product was excised from the agarose gel and then purified by QIAquick Gel Extraction Kit (Qiagen). Sequencing primers were those used for the RT-PCR analyses. Sequencing reaction was performed by means of the BigDye Terminator v1.1 Cycle Sequencing Kit (Applied Biosystems).

#### Quantitation of the relative level of correctly spliced transcripts in artificially introduced GT>GC mutations

The relative level of correctly spliced transcripts in association with GT>GC mutations that generated only wild-type transcripts (confirmed by Sanger sequencing) was determined by real-time quantitative RT-PCR analyses, essentially as described elsewhere [44, 46, 50]. Results were from three independent transfection experiments, with each experiment being performed in three replicates.

### Pictogram Analysis of the 9-bp 5’SS Signal Sequences Associated with 5’SS GT>GC mutations

The 9-bp canonical 5’SS signal sequences of the currently studied disease-associated and artificially introduced GT>GC mutations were extracted from the UCSC Genome Browser (https://genome.ucsc.edu/). The respective pictograms were constructed using WebLogo (http://weblogo.berkeley.edu/).

### *In Silico* Splicing Prediction

*In silico* splicing prediction was performed by means of Alamut^®^ Visual v.2.11 rev. 0 (https://www.interactive-biosoftware.com/; Interactive Biosoftware, Rouen, France) under default conditions.

## Supporting information

## Acknowledgements

We are grateful to the original authors who reported the disease-causing 5’SS GT>GC mutations studied here. We thank Nicolas Tomat and Léhna Bouchama (Brest, France) for technical assistance.

## REFERENCES

1. Turunen JJ, Niemela EH, Verma B, Frilander MJ. The significant other: splicing by the minor spliceosome. Wiley Interdiscip Rev RNA. 2013;4:61–76.

2. Parada GE, Munita R, Cerda CA, Gysling K. A comprehensive survey of non-canonical splice sites in the human transcriptome. Nucleic Acids Res. 2014;42:10564–78.

3. Verma B, Akinyi MV, Norppa AJ, Frilander MJ. Minor spliceosome and disease. Semin Cell Dev Biol. 2018;79:103–12.

4. Sharp PA, Burge CB. Classification of introns: U2-type or U12-type. Cell. 1997;91:875–9.

5. Papasaikas P, Valcarcel J. The spliceosome: the ultimate RNA chaperone and sculptor. Trends Biochem Sci. 2016;41:33–45.

6. Mount SM. A catalogue of splice junction sequences. Nucleic Acids Res. 1982;10:459–72.

7. Burset M, Seledtsov IA, Solovyev VV. Analysis of canonical and non-canonical splice sites in mammalian genomes. Nucleic Acids Res. 2000;28:4364–75.

8. Abril JF, Castelo R, Guigo R. Comparison of splice sites in mammals and chicken. Genome Res. 2005;15:111–9.

9. Roca X, Akerman M, Gaus H, Berdeja A, Bennett CF, Krainer AR. Widespread recognition of 5’ splice sites by noncanonical base-pairing to U1 snRNA involving bulged nucleotides. Genes Dev. 2012;26:1098–109.

10. Roca X, Krainer AR, Eperon IC. Pick one, but be quick: 5’ splice sites and the problems of too many choices. Genes Dev. 2013;27:129–44.

11. Wong MS, Kinney JB, Krainer AR. Quantitative activity profile and context dependence of all human 5’ splice sites. Mol Cell. 2018;71:1012–26 e3.

12. Mount SM, Pettersson I, Hinterberger M, Karmas A, Steitz JA. The U1 small nuclear RNA-protein complex selectively binds a 5’ splice site in vitro. Cell. 1983;33:509–18.

13. Kramer A, Keller W, Appel B, Luhrmann R. The 5’ terminus of the RNA moiety of U1 small nuclear ribonucleoprotein particles is required for the splicing of messenger RNA precursors. Cell. 1984;38:299–307.

14. Zhuang Y, Weiner AM. A compensatory base change in U1 snRNA suppresses a 5’ splice site mutation. Cell. 1986;46:827–35.

15. Kondo Y, Oubridge C, van Roon AM, Nagai K. Crystal structure of human U1 snRNP, a small nuclear ribonucleoprotein particle, reveals the mechanism of 5’ splice site recognition. Elife. 2015;4.

16. Dodgson JB, Engel JD. The nucleotide sequence of the adult chicken alpha-globin genes. J Biol Chem. 1983;258:4623–9.

17. Erbil C, Niessing J. The primary structure of the duck alpha D-globin gene: an unusual 5’ splice junction sequence. EMBO J. 1983;2:1339–43.

18. King CR, Piatigorsky J. Alternative RNA splicing of the murine alpha A-crystallin gene: protein-coding information within an intron. Cell. 1983;32:707–12.

19. Burset M, Seledtsov IA, Solovyev VV. SpliceDB: database of canonical and non-canonical mammalian splice sites. Nucleic Acids Res. 2001;29:255–9.

20. Sheth N, Roca X, Hastings ML, Roeder T, Krainer AR, Sachidanandam R. Comprehensive splice-site analysis using comparative genomics. Nucleic Acids Res. 2006;34:3955–67.

21. Churbanov A, Winters-Hilt S, Koonin EV, Rogozin IB. Accumulation of GC donor splice signals in mammals. Biol Direct. 2008;3:30.

22. Erkelenz S, Theiss S, Kaisers W, Ptok J, Walotka L, Muller L, et al. Ranking noncanonical 5’ splice site usage by genome-wide RNA-seq analysis and splicing reporter assays. Genome Res. 2018 Oct 24. doi:10.1101/gr.235861.118. [Epub ahead of print].

23. Aebi M, Hornig H, Padgett RA, Reiser J, Weissmann C. Sequence requirements for splicing of higher eukaryotic nuclear pre-mRNA. Cell. 1986;47:555–65.

24. Aebi M, Hornig H, Weissmann C. 5’ cleavage site in eukaryotic pre-mRNA splicing is determined by the overall 5’ splice region, not by the conserved 5’ GU. Cell. 1987;50:237–46.

25. Pagani F, Buratti E, Stuani C, Bendix R, Dork T, Baralle FE. A new type of mutation causes a splicing defect in ATM. Nat Genet. 2002;30:426–9.

26. Kralovicova J, Hwang G, Asplund AC, Churbanov A, Smith CI, Vorechovsky I. Compensatory signals associated with the activation of human GC 5’ splice sites. Nucleic Acids Res. 2011;39:7077–91.

27. Zhang XH, Arias MA, Ke S, Chasin LA. Splicing of designer exons reveals unexpected complexity in pre-mRNA splicing. RNA. 2009;15:367–76.

28. De Conti L, Baralle M, Buratti E. Exon and intron definition in pre-mRNA splicing. Wiley Interdiscip Rev RNA. 2013;4:49–60.

29. Boehm V, Britto-Borges T, Steckelberg AL, Singh KK, Gerbracht JV, Gueney E, et al. Exon junction complexes suppress spurious splice sites to safeguard rranscriptome integrity. Mol Cell. 2018;72:482–95 e7.

30. Ramalho AS, Beck S, Meyer M, Penque D, Cutting GR, Amaral MD. Five percent of normal cystic fibrosis transmembrane conductance regulator mRNA ameliorates the severity of pulmonary disease in cystic fibrosis. Am J Respir Cell Mol Biol. 2002;27:619–27.

31. Raraigh KS, Han ST, Davis E, Evans TA, Pellicore MJ, McCague AF, et al. Functional assays are essential for interpretation of missense variants associated with variable expressivity. Am J Hum Genet. 2018;102:1062–77.

32. Stenson PD, Mort M, Ball EV, Evans K, Hayden M, Heywood S, et al. The Human Gene Mutation Database: towards a comprehensive repository of inherited mutation data for medical research, genetic diagnosis and next-generation sequencing studies. Hum Genet. 2017;136:665–77.

33. Desviat LR, Clavero S, Perez-Cerda C, Navarrete R, Ugarte M, Perez B. New splicing mutations in propionic acidemia. J Hum Genet. 2006;51:992–7.

34. Soudais C, de Villartay JP, Le Deist F, Fischer A, Lisowska-Grospierre B. Independent mutations of the human CD3-epsilon gene resulting in a T cell receptor/CD3 complex immunodeficiency. Nat Genet. 1993;3:77–81.

35. Lin P, Li W, Wen B, Zhao Y, Fenster DS, Wang Y, et al. Novel *PNPLA2* gene mutations in Chinese Han patients causing neutral lipid storage disease with myopathy. J Hum Genet. 2012;57:679– 81.

36. Seyama K, Nonoyama S, Gangsaas I, Hollenbaugh D, Pabst HF, Aruffo A, et al. Mutations of the CD40 ligand gene and its effect on CD40 ligand expression in patients with X-linked hyper IgM syndrome. Blood. 1998;92:2421–34.

37. Bartolo C, Papp AC, Snyder PJ, Sedra MS, Burghes AH, Hall CD, et al. A novel splice site mutation in a Becker muscular dystrophy patient. J Med Genet. 1996;33:324–7.

38. Hastbacka J, Kerrebrock A, Mokkala K, Clines G, Lovett M, Kaitila I, et al. Identification of the Finnish founder mutation for diastrophic dysplasia (DTD). Eur J Hum Genet. 1999;7:664–70.

39. Muller JS, Piko H, Schoser BG, Schlotter-Weigel B, Reilich P, Gurster S, et al. Novel splice site mutation in the caveolin-3 gene leading to autosomal recessive limb girdle muscular dystrophy. Neuromuscul Disord. 2006;16:432–6.

40. Aoyagi Y, Kobayashi H, Tanaka K, Ozawa T, Nitta H, Tsuji S. A de novo splice donor site mutation causes in-frame deletion of 14 amino acids in the proteolipid protein in Pelizaeus-Merzbacher disease. Ann Neurol. 1999;46:112–5.

41. Kume K, Masamune A, Kikuta K, Shimosegawa T. [−215G>A; IVS3+2T>C] mutation in the *SPINK1* gene causes exon 3 skipping and loss of the trypsin binding site. Gut. 2006;55:1214.

42. Zou WB, Boulling A, Masson E, Cooper DN, Liao Z, Li ZS, et al. Clarifying the clinical relevance of *SPINK1* intronic variants in chronic pancreatitis. Gut. 2016;65:884–6.

43. Zou WB, Masson E, Boulling A, Cooper DN, Li ZS, Liao Z, et al. Digging deeper into the intronic sequences of the *SPINK1* gene. Gut. 2016;65:1055–6.

44. Boulling A, Chen JMC, I., Férec C. Is the SPINK1 p.Asn34Ser missense mutation per se the true culprit within its associated haplotype? WebmedCentral GENETICS. 2012;3:WMC003084 (Available at: https://www.webmedcentral.com/article_view/3084). Accessed 14 November 2018.

45. Fu XD, Ares M, Jr. Context-dependent control of alternative splicing by RNA-binding proteins. Nat Rev Genet. 2014;15:689–701.

46. Zou WB, Boulling A, Masamune A, Issarapu P, Masson E, Wu H, et al. No association between *CEL-HYB* hybrid allele and chronic pancreatitis in Asian populations. Gastroenterology. 2016;150:1558–60 e5.

47. Lykke-Andersen S, Jensen TH. Nonsense-mediated mRNA decay: an intricate machinery that shapes transcriptomes. Nat Rev Mol Cell Biol. 2015;16:665–77.

48. Popp MW, Maquat LE. Leveraging rules of nonsense-mediated mRNA decay for genome engineering and personalized medicine. Cell. 2016;165:1319–22.

49. Pereverzev AP, Gurskaya NG, Ermakova GV, Kudryavtseva EI, Markina NM, Kotlobay AA, et al. Method for quantitative analysis of nonsense-mediated mRNA decay at the single cell level. Sci Rep. 2015;5:7729.

50. Zou WB, Wu H, Boulling A, Cooper DN, Li ZS, Liao Z, et al. *In silico* prioritization and further functional characterization of *SPINK1* intronic variants. Hum Genomics. 2017;11:7.

51. Wu H, Boulling A, Cooper DN, Li ZS, Liao Z, Chen JM, et al. *In vitro* and *in silico* evidence against a significant effect of the *SPINK1* c.194G>A variant on pre-mRNA splicing. Gut. 2017;66:2195–6.

52. Ota Y, Masamune A, Inui K, Kume K, Shimosegawa T, Kikuyama M. Phenotypic variability of the homozygous IVS3+2T>C mutation in the serine protease inhibitor Kazal type 1 (*SPINK1*) gene in patients with chronic pancreatitis. Tohoku J Exp Med. 2010;221:197–201.

53. Venet T, Masson E, Talbotec C, Billiemaz K, Touraine R, Gay C, et al. Severe infantile isolated exocrine pancreatic insufficiency caused by the complete functional loss of the *SPINK1* gene. Hum Mutat. 2017;38:1660–5.

54. Yilmaz D, Cogulu O, Ozkinay F, Kavakli K, Roos D. A novel mutation in the *DIA1* gene in a patient with methemoglobinemia type II. Am J Med Genet A. 2005;133A:101-2.

55. Frischknecht H, Dutly F, Walker L, Nakamura-Garrett LM, Eng B, Waye JS. Three new beta-thalassemia mutations with varying degrees of severity. Hemoglobin. 2009;33:220–5.

56. Hladnik U, Nyhan WL, Bertelli M. Variable expression of HPRT deficiency in 5 members of a family with the same mutation. Arch Neurol. 2008;65:1240–3.

57. Wuhl E, Kogan J, Zurowska A, Matejas V, Vandevoorde RG, Aigner T, et al. Neurodevelopmental deficits in Pierson (microcoria-congenital nephrosis) syndrome. Am J Med Genet A. 2007;143:311–9.

58. Bar DZ, Arlt MF, Brazier JF, Norris WE, Campbell SE, Chines P, et al. A novel somatic mutation achieves partial rescue in a child with Hutchinson-Gilford progeria syndrome. J Med Genet. 2017;54:212–6.

59. Al-Mahdili HA, Hooper AJ, Sullivan DR, Stewart PM, Burnett JR. A mild case of abetalipoproteinaemia in association with subclinical hypothyroidism. Ann Clin Biochem. 2006;43:516–9.

60. Kajihara S, Hisatomi A, Mizuta T, Hara T, Ozaki I, Wada I, et al. A splice mutation in the human canalicular multispecific organic anion transporter gene causes Dubin-Johnson syndrome. Biochem Biophys Res Commun. 1998;253:454–7.

61. Fukao T, Yamaguchi S, Scriver CR, Dunbar G, Wakazono A, Kano M, et al. Molecular studies of mitochondrial acetoacetyl-coenzyme A thiolase deficiency in the two original families. Hum Mutat. 1993;2:214–20.

62. Lagier-Tourenne C, Tazir M, Lopez LC, Quinzii CM, Assoum M, Drouot N, et al. ADCK3, an ancestral kinase, is mutated in a form of recessive ataxia associated with coenzyme Q10 deficiency. Am J Hum Genet. 2008;82:661–72.

63. Dolcini L, Caridi G, Dagnino M, Sala A, Gokce S, Sokucu S, et al. Analbuminemia produced by a novel splicing mutation. Clin Chem. 2007;53:1549–52.

64. Infante JB, Alvelos MI, Bastos M, Carrilho F, Lemos MC. Complete androgen insensitivity syndrome caused by a novel splice donor site mutation and activation of a cryptic splice donor site in the androgen receptor gene. J Steroid Biochem Mol Biol. 2016;155:63–6.

65. Rios M, Storry JR, Hue-Roye K, Chung A, Reid ME. Two new molecular bases for the Dombrock null phenotype. Br J Haematol. 2002;117:765–7.

66. Das S, Levinson B, Whitney S, Vulpe C, Packman S, Gitschier J. Diverse mutations in patients with Menkes disease often lead to exon skipping. Am J Hum Genet. 1994;55:883–9.

67. Hopp K, Heyer CM, Hommerding CJ, Henke SA, Sundsbak JL, Patel S, et al. *B9D1* is revealed as a novel Meckel syndrome (MKS) gene by targeted exon-enriched next-generation sequencing and deletion analysis. Hum Mol Genet. 2011;20:2524–34.

68. Haire RN, Ohta Y, Strong SJ, Litman RT, Liu Y, Prchal JT, et al. Unusual patterns of exon skipping in Bruton tyrosine kinase are associated with mutations involving the intron 17 3’ splice site. Am J Hum Genet. 1997;60:798–807.

69. Rahner N, Nuernberg G, Finis D, Nuernberg P, Royer-Pokora B. A novel *C8orf37* splice mutation and genotype-phenotype correlation for cone-rod dystrophy. Ophthalmic Genet. 2016;37:294– 300.

70. Tosetto E, Ghiggeri GM, Emma F, Barbano G, Carrea A, Vezzoli G, et al. Phenotypic and genetic heterogeneity in Dent’s disease-the results of an Italian collaborative study. Nephrol Dial Transplant. 2006;21:2452–63.

71. Nicholls AC, Valler D, Wallis S, Pope FM. Homozygosity for a splice site mutation of the *COL1A2* gene yields a non-functional pro(alpha)2(I) chain and an EDS/OI clinical phenotype. J Med Genet. 2001;38:132–6.

72. Haas JT, Winter HS, Lim E, Kirby A, Blumenstiel B, DeFelice M, et al. *DGAT1* mutation is linked to a congenital diarrheal disorder. J Clin Invest. 2012;122:4680–4.

73. Wibawa T, Takeshima Y, Mitsuyoshi I, Wada H, Surono A, Nakamura H, et al. Complete skipping of exon 66 due to novel mutations of the dystrophin gene was identified in two Japanese families of Duchenne muscular dystrophy with severe mental retardation. Brain Dev. 2000;22:107–12.

74. Ahmed I, Mittal K, Sheikh TI, Vasli N, Rafiq MA, Mikhailov A, et al. Identification of a homozygous splice site mutation in the dynein axonemal light chain 4 gene on 22q13.1 in a large consanguineous family from Pakistan with congenital mirror movement disorder. Hum Genet. 2014;133:1419–29.

75. Hermans MM, van Leenen D, Kroos MA, Reuser AJ. Mutation detection in glycogen storage-disease type II by RT-PCR and automated sequencing. Biochem Biophys Res Commun. 1997;241:414–8.

76. Sobrier ML, Maghnie M, Vie-Luton MP, Secco A, di Iorgi N, Lorini R, et al. Novel *HESX1* mutations associated with a life-threatening neonatal phenotype, pituitary aplasia, but normally located posterior pituitary and no optic nerve abnormalities. J Clin Endocrinol Metab. 2006;91:4528–36.

77. Moran CJ, Walters TD, Guo CH, Kugathasan S, Klein C, Turner D, et al. IL-10R polymorphisms are associated with very-early-onset ulcerative colitis. Inflamm Bowel Dis. 2013;19:115–23.

78. Humbert C, Silbermann F, Morar B, Parisot M, Zarhrate M, Masson C, et al. Integrin alpha 8 recessive mutations are responsible for bilateral renal agenesis in humans. Am J Hum Genet. 2014;94:288–94.

79. Vockley J, Rogan PK, Anderson BD, Willard J, Seelan RS, Smith DI, et al. Exon skipping in *IVD* RNA processing in isovaleric acidemia caused by point mutations in the coding region of the *IVD* gene. Am J Hum Genet. 2000;66:356–67.

80. Villa A, Sironi M, Macchi P, Matteucci C, Notarangelo LD, Vezzoni P, et al. Monocyte function in a severe combined immunodeficient patient with a donor splice site mutation in the *Jak3* gene. Blood. 1996;88:817–23.

81. Allamand V, Sunada Y, Salih MA, Straub V, Ozo CO, Al-Turaiki MH, et al. Mild congenital muscular dystrophy in two patients with an internally deleted laminin alpha2-chain. Hum Mol Genet. 1997;6:747–52.

82. Nichols WC, Seligsohn U, Zivelin A, Terry VH, Hertel CE, Wheatley MA, et al. Mutations in the ER-Golgi intermediate compartment protein ERGIC-53 cause combined deficiency of coagulation factors V and VIII. Cell. 1998;93:61–70.

83. Wood-Trageser MA, Gurbuz F, Yatsenko SA, Jeffries EP, Kotan LD, Surti U, et al. *MCM9* mutations are associated with ovarian failure, short stature, and chromosomal instability. Am J Hum Genet. 2014;95:754–62.

84. Gok F, Crettol LM, Alanay Y, Hacihamdioglu B, Kocaoglu M, Bonafe L, et al. Clinical and radiographic findings in two brothers affected with a novel mutation in matrix metalloproteinase 2 gene. Eur J Pediatr. 2010;169:363–7.

85. Martin CA, Murray JE, Carroll P, Leitch A, Mackenzie KJ, Halachev M, et al. Mutations in genes encoding condensin complex proteins cause microcephaly through decatenation failure at mitosis. Genes Dev. 2016;30:2158–72.

86. Tanugi-Cholley LC, Issartel JP, Lunardi J, Freycon F, Morel F, Vignais PV. A mutation located at the 5’ splice junction sequence of intron 3 in the p67phox gene causes the lack of p67phox mRNA in a patient with chronic granulomatous disease. Blood. 1995;85:242–9.

87. Zanni G, Saillour Y, Nagara M, Billuart P, Castelnau L, Moraine C, et al. Oligophrenin 1 mutations frequently cause X-linked mental retardation with cerebellar hypoplasia. Neurology. 2005;65:1364–9.

88. Matsuura T, Hoshide R, Komaki S, Kiwaki K, Endo F, Nakamura S, et al. Identification of two new aberrant splicings in the ornithine carbamoyltransferase (*OCT*) gene in two patients with early and late onset OCT deficiency. J Inherit Metab Dis. 1995;18:273–82.

89. Shimozawa N, Nagase T, Takemoto Y, Suzuki Y, Fujiki Y, Wanders RJ, et al. A novel aberrant splicing mutation of the *PEX16* gene in two patients with Zellweger syndrome. Biochem Biophys Res Commun. 2002;292:109–12.

90. Biancheri R, Grossi S, Regis S, Rossi A, Corsolini F, Rossi DP, et al. Further genotype-phenotype correlation emerging from two families with *PLP1* exon 4 skipping. Clin Genet. 2014;85:267–72.

91. Aldahmesh MA, Mohamed JY, Alkuraya FS. A novel mutation in *PRDM5* in brittle cornea syndrome. Clin Genet. 2012;81:198–9.

92. Smith SB, Qu HQ, Taleb N, Kishimoto NY, Scheel DW, Lu Y, et al. Rfx6 directs islet formation and insulin production in mice and humans. Nature. 2010;463:775–80.

93. Adly N, Alhashem A, Ammari A, Alkuraya FS. Ciliary genes *TBC1D32/C6orf170* and *SCLT1* are mutated in patients with OFD type IX. Hum Mutat. 2014;35:36–40.

94. Sumegi J, Huang D, Lanyi A, Davis JD, Seemayer TA, Maeda A, et al. Correlation of mutations of the *SH2D1A* gene and epstein-barr virus infection with clinical phenotype and outcome in X-linked lymphoproliferative disease. Blood. 2000;96:3118–25.

95. Leman R, Gaildrat P, Gac GL, Ka C, Fichou Y, Audrezet MP, et al. Novel diagnostic tool for prediction of variant spliceogenicity derived from a set of 395 combined in silico/in vitro studies: an international collaborative effort. Nucleic Acids Res. 2018;46:7913–23.

