## Supplementary material for "First estimation of the scale of canonical 5’ splice site GT>GC mutations generating wild-type transcripts and their medical genetic implications"

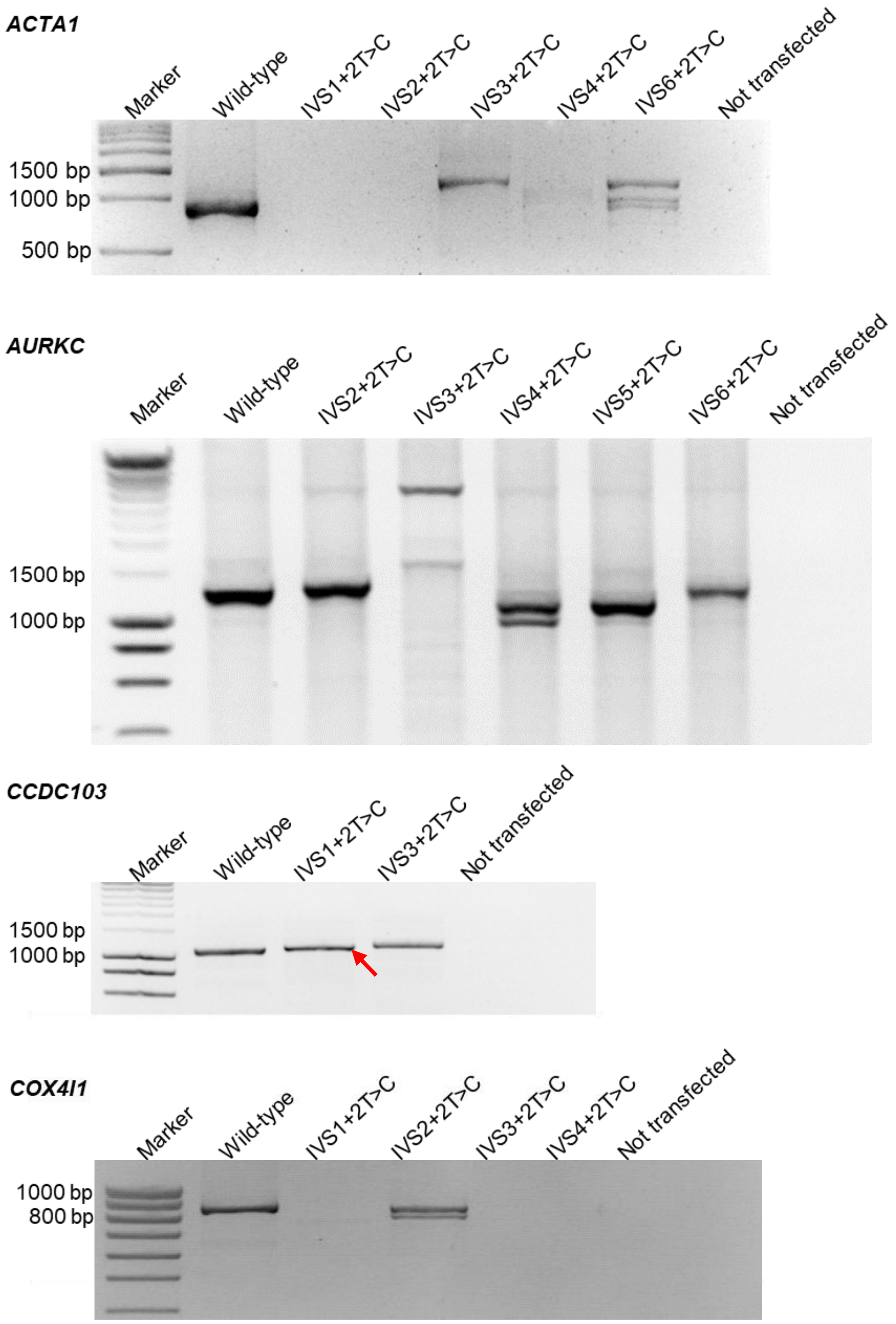


**Supplementary Figure S1.** RT-PCR results from all the currently analyzed 5'SS GT>GC mutations by means of full-length gene splicing assay. Wild-type transcripts (confirmed by sequencing) resulting from the +2T>C mutations are indicated by arrows. IVS, InterVening Sequence (i.e., an intron).

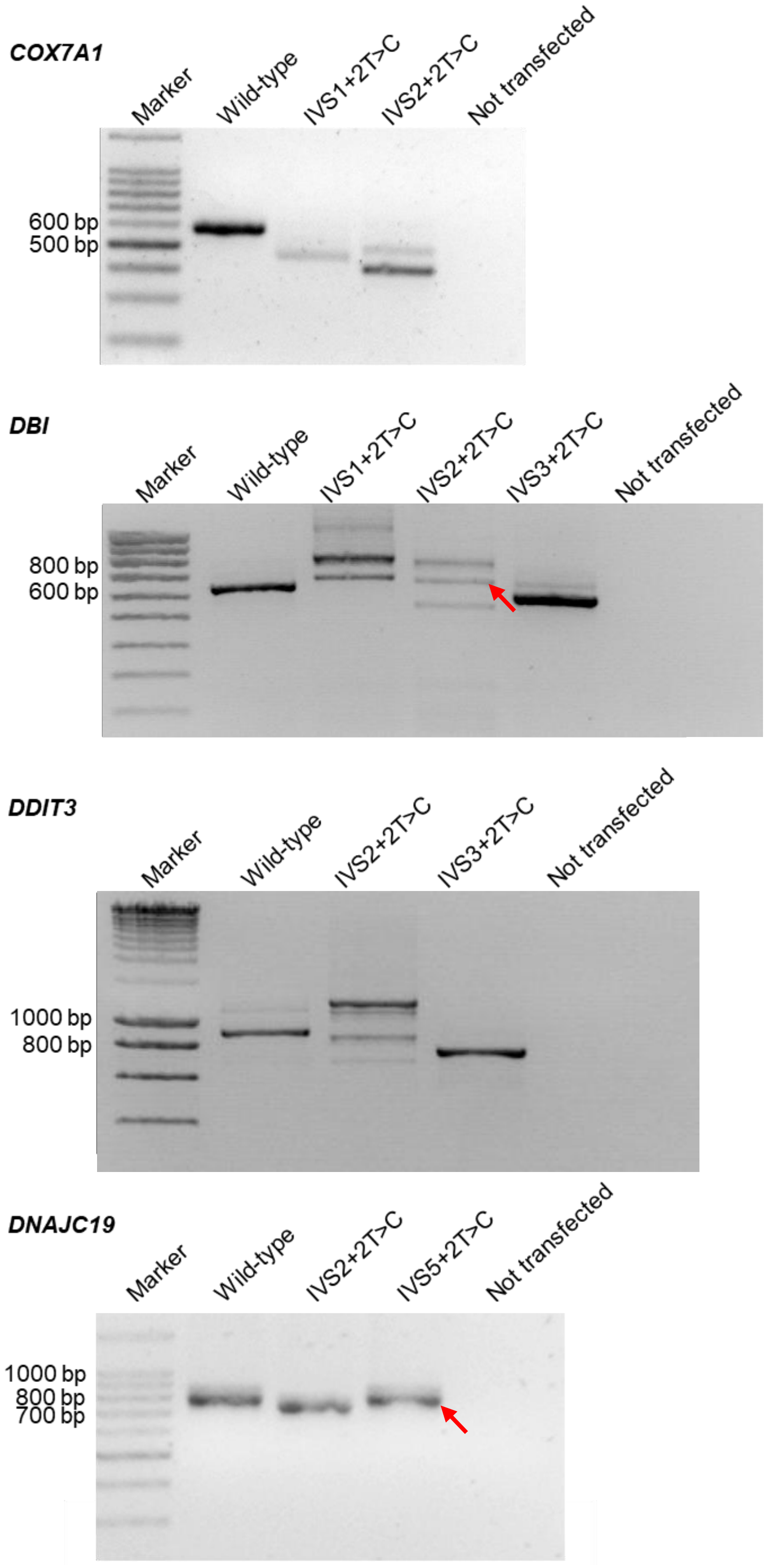


**Figure S1** (*continued*)

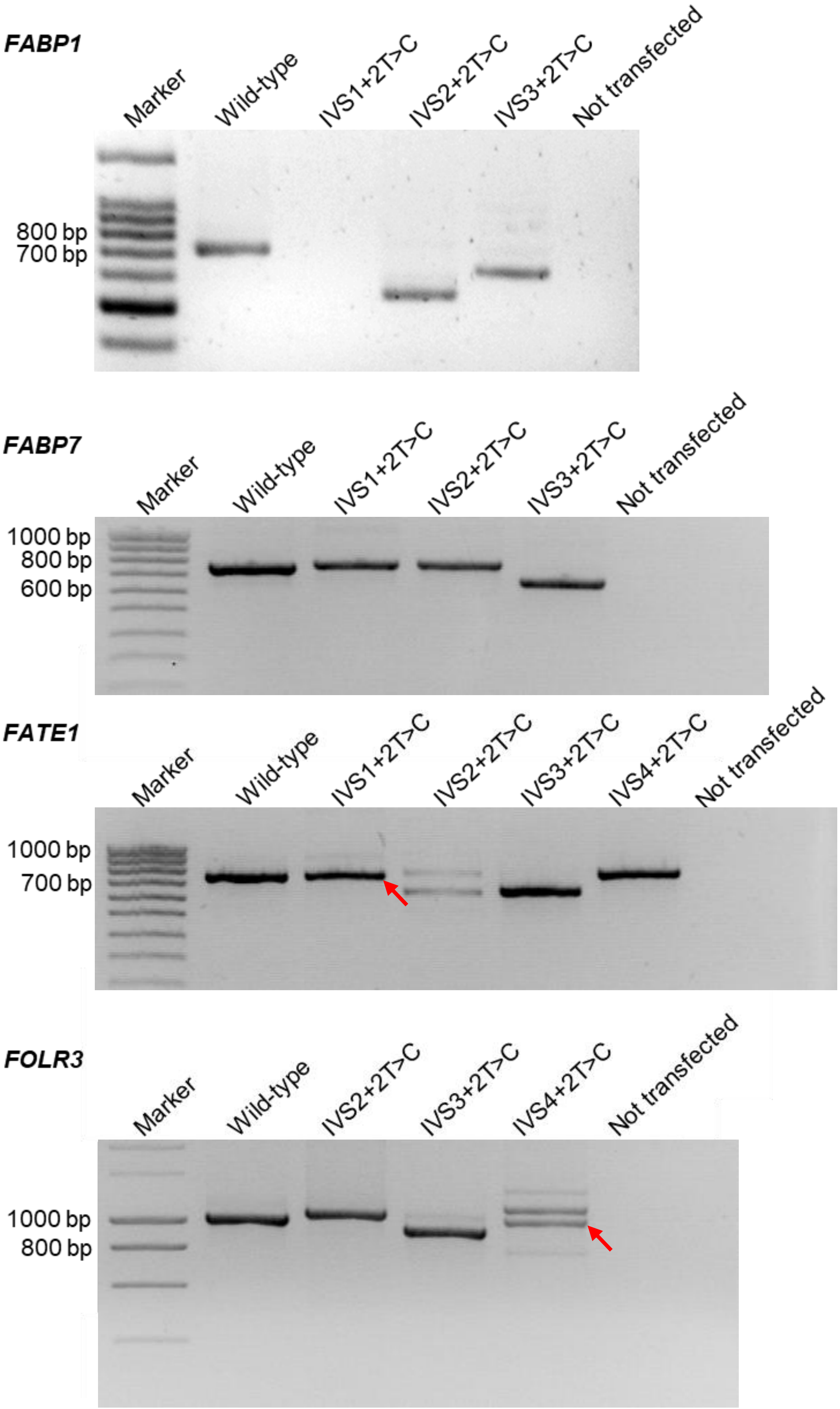


**Figure S1** (*continued*)

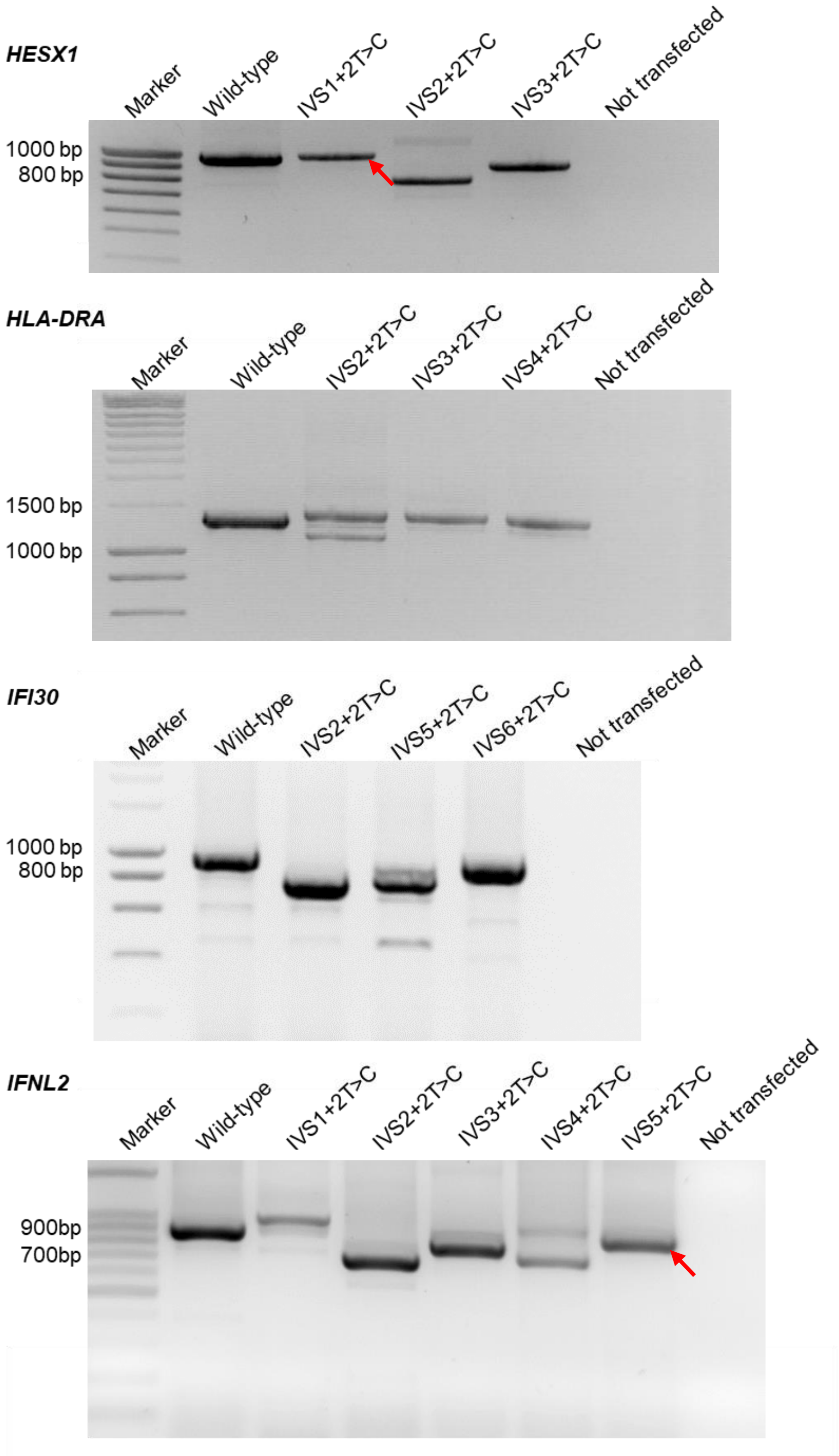


**Figure S1** (*continued*)

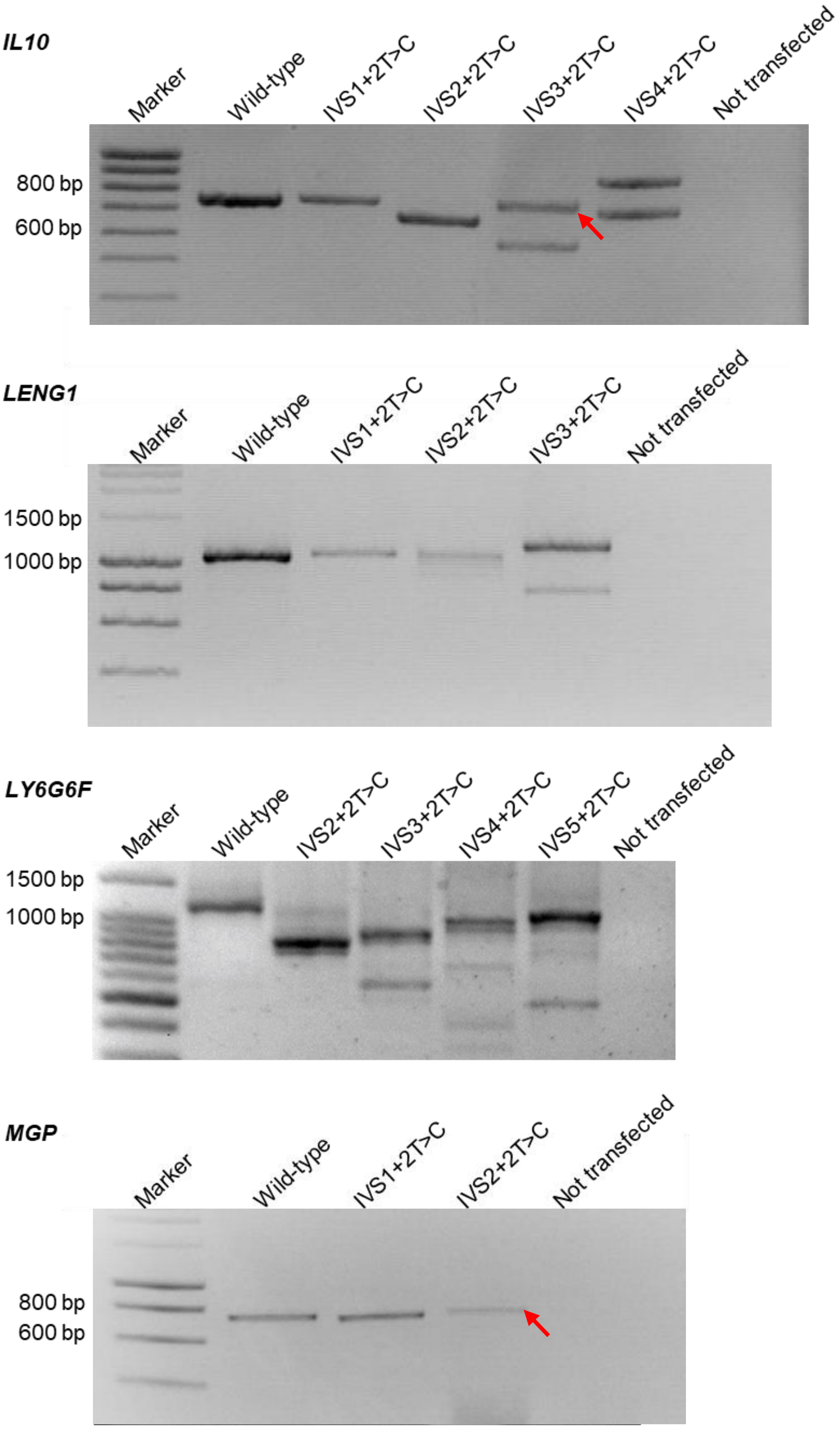


**Figure S1** (*continued*)

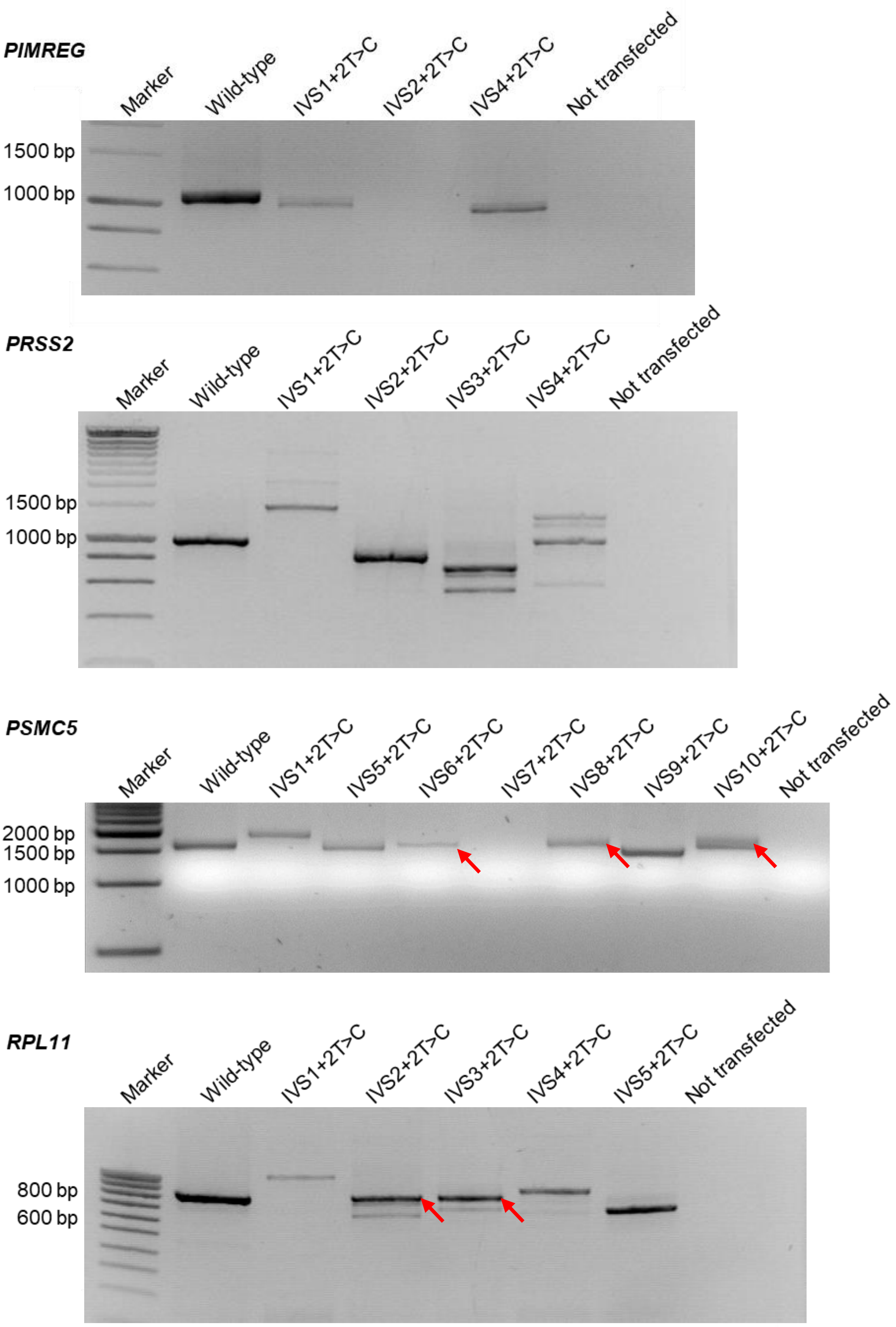


**Figure S1** (*continued*)

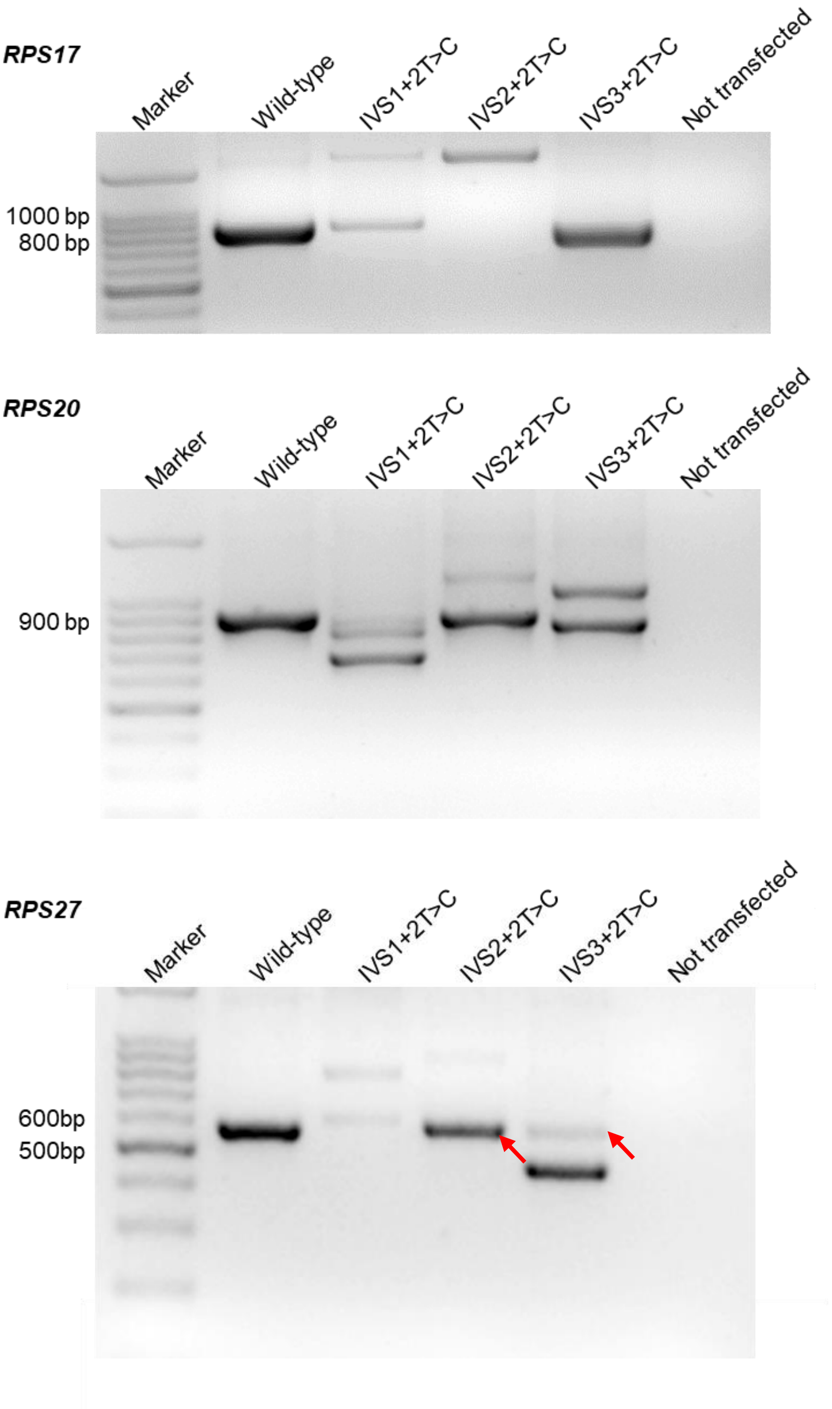


**Figure S1** (*continued*)

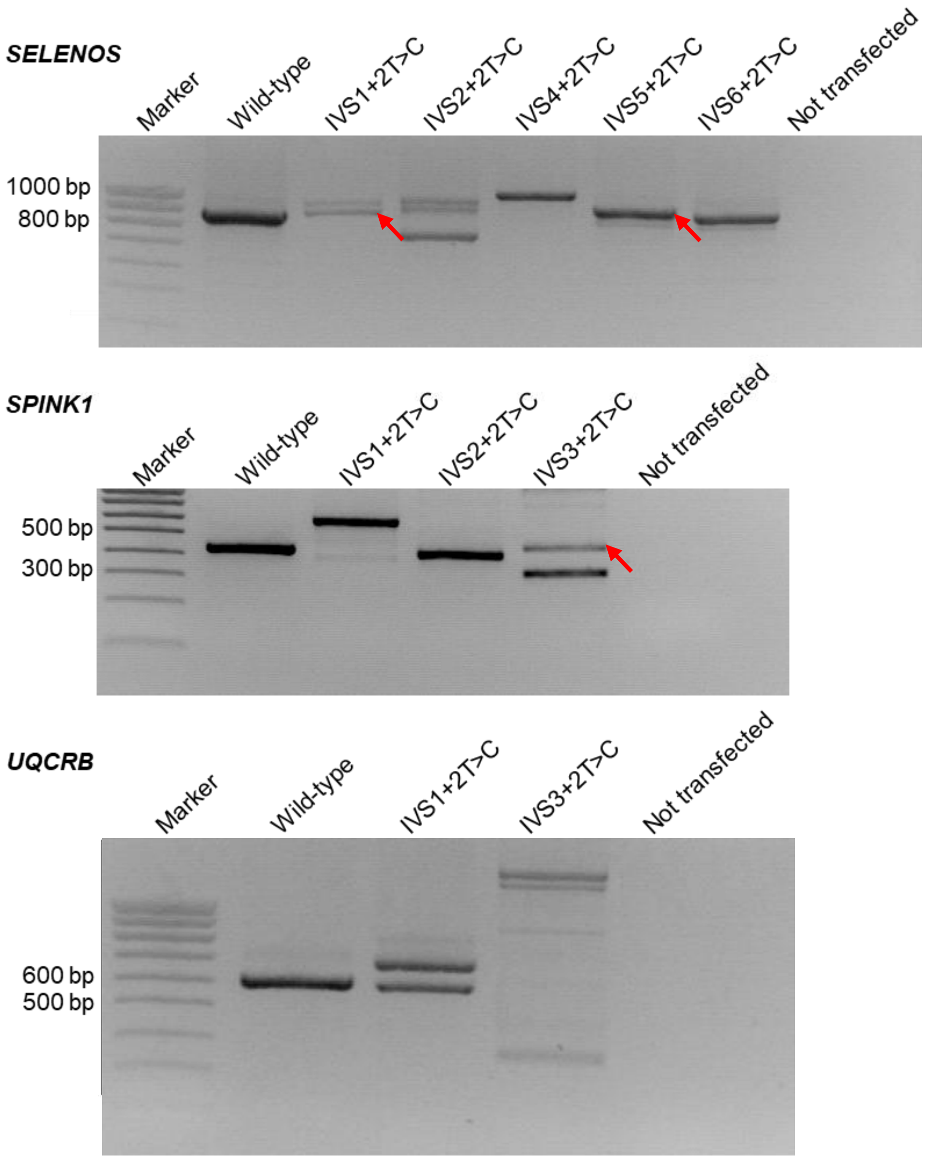


**Figure S1** (*continued*)

**a**

5'......CGTGGTCTGCGCGGGGTTGCCCTCCTGTTCTGGTTTATCAGGGGATCCCCAAAGAAAGCAAGGGGACCAAGGCCGG
GACTGCTGGGGTGAAG**GT**CCGGGAGGCTGAGTAAGGGGACGGAAGG**gt**tagttcta[...]gccttccc**ag**GCACAGGCCATGGAAAGG
AATGACATCATCAACTTCAAGGCTTTGGAGAAAGAGCTGCAGGCTGCACTCACTGCTGATGAGAAGTACAAACGGGAG......3'

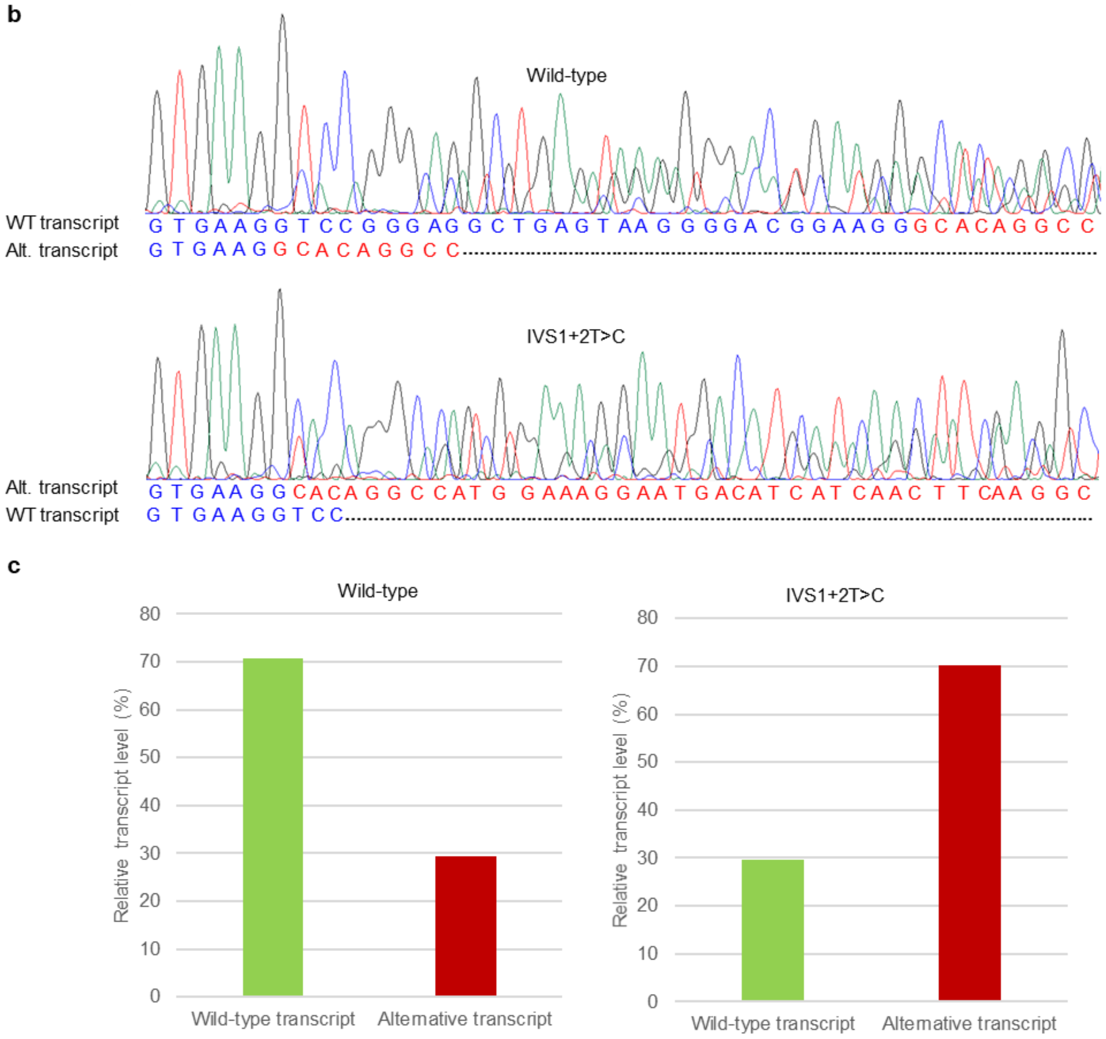


**Supplementary Figure S2.** Rough estimation of the relative expression level of the *CCDC103* IVS1+2T>C allele-derived wild-type transcript compared with that derived from the wild-type allele. (**A**) Illustration of partial genomic sequence of the *CCDC103* gene. The normally spliced exons 1 and 2 are shown in blue and red capital letters, the corresponding intron being shown in lower case letters and the canonical GT-AG splice sites being highlighted in bold. The cryptic 5'SS GT (located within exon 1) used for an alternatively spliced transcript is in bold and underlined. The two transcripts were indistinguishable in RT-PCR analysis (refer to Supplementary Figure S1). (**B**) Sequencing electropherograms showing the co-existence of the wild-type (WT) transcript and the alternative (alt.) transcript in the RT-PCR products from both the wild-type *CCDC103* gene and the *CCDC103* IVS1+2T>C mutant. (**C**) Relative expression levels of the two transcripts generated from the wild-type *CCDC103* gene and the *CCDC103* IVS1+2T>C mutant. The level of the correctly spliced transcripts generated from the mutant allele was calculated to be ~18% [(30% × 30%)/(70% × 70%)] of that generated from the wild-type allele.

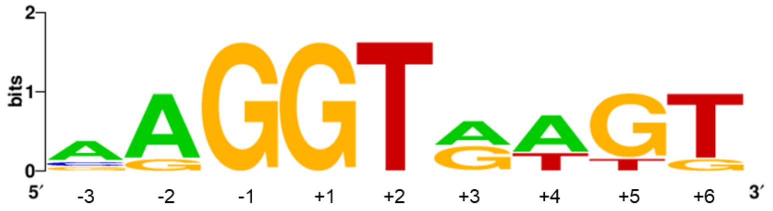


**Supplementary Figure S3.** Pictogram of the 6 5'SSs containing GT>GC mutations previously reported to confer a milder than expected phenotype but having no supportive patient-derived transcript expression data.
